# Lateral frontoparietal effective connectivity differentiates and predicts state of consciousness in traumatic disorders of consciousness

**DOI:** 10.1101/2023.06.07.544105

**Authors:** Riku Ihalainen, Jitka Annen, Olivia Gosseries, Paolo Cardone, Rajanikant Panda, Charlotte Martial, Aurore Thibaut, Steven Laureys, Srivas Chennu

## Abstract

Neuroimaging studies have suggested an important role for the default mode network (DMN) in disorders of consciousness (DoC). However, the extent to which DMN connectivity can discriminate DoC states – unresponsive wakefulness syndrome (UWS) and minimally conscious state (MCS) – is less evident. Particularly, it is unclear whether effective DMN connectivity, as measured indirectly with dynamic causal modelling (DCM) of resting EEG can disentangle UWS from healthy controls and from patients considered conscious (MCS+). Crucially, this extends to UWS patients with potentially “covert” awareness (minimally conscious star, MCS*) indexed by voluntary brain activity in conjunction with partially preserved frontoparietal metabolism as measured with positron emission tomography (PET+ diagnosis; in contrast to PET-diagnosis with complete frontoparietal hypometabolism). Here, we address this gap by using DCM of EEG data acquired from patients with traumatic brain injury in 11 UWS (6 PET– and 5 PET+) and in 12 MCS+ (11 PET+ and 1 PET-), alongside with 11 healthy controls. We provide evidence for a key difference in left frontoparietal connectivity when contrasting UWS PET– with MCS+ patients and healthy controls. Next, in a leave-one-subject-out cross-validation, we tested the classification performance of the DCM models demonstrating that connectivity between medial prefrontal and left parietal sources reliably discriminates UWS PET– from MCS+ patients and controls. Finally, we illustrate that these models generalize to an unseen dataset: models trained to discriminate UWS PET– from MCS+ and controls, classify MCS* patients as conscious subjects with high posterior probability (pp > .92). These results identify specific alterations in the DMN after severe brain injury and highlight the clinical utility of EEG– based effective connectivity for identifying patients with potential covert awareness.

**Author Summary:** Our study investigates the role of the Default Mode Network (DMN) in individuals with disorders of consciousness (DoC), such as unresponsive wakefulness syndrome (UWS) and minimally conscious state (MCS). Previous neuroimaging studies have suggested a role for the DMN in DoC, but its ability to differentiate between UWS and MCS remain unclear.

Using advance brain imaging and modelling techniques, we analyzed data from DoC patients with traumatic brain injury and healthy controls. Our findings reveal a key difference in left frontoparietal connectivity when comparing UWS to MCS patients and healthy individuals.

To validate our results, we employed a robust cross-validation approach, which demonstrated that the connectivity between frontal and left parietal brain regions reliably discriminates UWS patients from MCS patients and controls. Furthermore, we extended our analysis to include patients with potential covert awareness, showcasing the clinical utility of our findings. We successfully classified these patients as conscious with high accuracy.

This research significantly contributes to our understanding of the DMN in DoC and highlights the potential use of electroencephalography-based connectivity analysis in clinical settings. By identifying specific alterations in the DMN after severe brain injury, our study may aid in the accurate diagnosis and management of individuals with disorders of consciousness, potentially improving their overall outcomes.

## 1 Introduction

After a severe brain injury, patients may be diagnosed with a transient or permanent disorders of consciousness (DoC), such as the unresponsive wakefulness syndrome (UWS) or the minimally conscious state (MCS). The UWS is defined by preserved arousal in the absence of behavioral signs of awareness (periodic sustained eye opening with purposeless movements; Laureys et al., 2010). In contrast, patients in MCS show fluctuating but reproducible signs of consciousness with preserved arousal. The MCS has been further divided into MCS– and MCS+, with the latter condition characterized by command following, intelligible verbalization, or gestural or verbal yes/no responses (regardless of accuracy) to spoken or written questions (Bruno et al., 2011).

The exclusive use of clinical consensus for diagnosing these DoC based on observed behaviors has been shown to result in high rates of misdiagnosis of the accurate level of consciousness of the DoC patients, especially in the case of patients suffering from UWS (Stender et al., 2014; Thibaut et al., 2021; van Erp et al., 2015). Consequently, with the advent of modern neuroimaging techniques, there has been increasing interest in characterizing the underlying neuronal basis for the presence or lack of awareness in DoC using structural and functional magnetic resonance imaging (MRI/fMRI; e.g., Demertzi et al., 2015; Di Perri et al., 2016), positron emission tomography (PET; e.g., Laureys et al., 1999; Stender et al., 2014), and electroencephalography (EEG; e.g., Chennu et al., 2014; King et al., 2013; Sitt et al., 2014).

Structural and functional neuroimaging studies have suggested an important role of the default mode network (DMN) in DoC – an intrinsic brain network encompassing the posterior cingulate cortex/precuneus, bilateral parietal cortices, and the medial prefrontal cortex (Annen et al., 2018; Boly et al., 2009; Fernández-Espejo et al., 2012; Guldenmund et al., 2016; Soddu et al., 2012; Vanhaudenhuyse et al., 2010). In parallel, cerebral metabolism as measured by PET has been shown to differentiate UWS from MCS (Stender et al., 2014; Stender et al., 2016; Thibaut et al., 2021), with regional differences often in areas associated with DMN (Stender et al., 2015; Thibaut et al., 2012). This extends to patients with MCS; MCS+ can be distinguished from MCS– with the former group showing partially preserved language related behaviors (e.g., response to simple commands) alongside with a higher cerebral metabolism especially in left-sided cortical areas, including Broca’s and Wernicke’s areas, premotor, presupplementary motor, and sensorimotor cortices (Aubinet et al., 2020; Bruno et al., 2012; Thibaut et al., 2020). A trained neurologist can diagnose patients also based on a visual inspection of their underlying PET metabolism, to as either PET negative (-) or PET positive (+). A PET-diagnosis is typically produced by a complete bilateral hypometabolism of the associative frontoparietal cortex with no voxels with preserved metabolism, whereas PET+ diagnosis is characterized by an incomplete hypometabolism and partial preservation of activity within these areas (Laureys et al., 2004; Thibaut et al., 2012, Stender et al. 2014).

In addition, effective connectivity studies in DoC as measured with dynamic causal modelling (DCM) for fMRI have suggested disruptions within the DMN specifically related to posterior cingulate cortex (PCC; Crone et al., 2015) and in subcortical networks, potentially driving the disruptions in the DMN (Chen et al., 2018; Coulborn et al., 2021). DCM is a generic approach used to infer hidden (or unobserved) neuronal states from measured brain activity; the idea is to model the source activity over time in terms of causal relationships between interacting inhibitory and excitatory populations of neurons. As far as we know, only one study has used DCM with EEG for measuring and diagnosing cognitive functioning in DoC population. Using a mismatch negativity paradigm, Boly and colleagues (2011a) showed that the difference between UWS and MCS was due to an impairment of backward connectivity from frontal to temporal cortices, emphasizing the importance of top-down processing for conscious perception.

Importantly, a number of studies have suggested residual consciousness and/or reported “covert” voluntary brain activity in some seemingly unresponsive patients, with both, active and resting state paradigms (Bodart et al., 2017; Claassen et al., 2019; Chennu et al., 2017; Cruse et al., 2011; Lechinger et al., 2013; Monti et al., 2010; Owen et al., 2006; Owen & Coleman, 2008; Schnakers et al., 2015). These patients, who show no behavioral signs of consciousness, yet with whose neuroimaging results indicate residual brain activity compatible with the diagnosis of MCS, have been termed MCS* (minimally conscious state star; Gosseries et al., 2014; Thibaut et al., 2021). To keep consistent with the literature, from this point on, we use the term MCS* to refer to the UWS patients with PET+ diagnosis in this manuscript.

Currently it is unknown whether effective resting state connectivity between key nodes within the DMN, as measured with EEG, could be used to identify such covertly aware patients. Here, as a preliminary investigation, we address this gap by using spectral DCM for EEG with parametric empirical Bayes (PEB). We investigate the difference in causal interactions between cortico-cortical regions of the DMN, between DoC patients (UWS and MCS+) and healthy controls. First, our interest is in distinguishing the differences between UWS patients and healthy controls, and in demonstrating the prospective performance of the connectivity within DMN in classifying these states. Crucially, we include MCS+ patients to function as a second, yet demonstrably conscious, control group to reduce the probability that our findings reflect mainly damage in the brain, and not consciousness itself. Based on previous studies (Boly et al., 2011a), we hypothesize that there will be top-down/backward connectivity differences in UWS vs. healthy controls and in UWS vs. MCS+ comparisons. We also model the difference between MCS+ and healthy controls where we do not expect to see this difference.

Next, in a leave-one-subject-out cross-validation, we test the classification performance of models based on the fully connected DMN network and on two connectivity subsets of the DMN: the posterior connections and the frontoparietal connections. Following this, we adopt a data-driven approach to the classification problem by investigating the predictive performance of single connections. The aim here is to identify the direction and location of the largest, most consistent modulations between the subjects.

Finally, we demonstrate that our DCM models generalize to a more difficult classification problem: in a leave-one-*state*-out cross-validation paradigm, we train the models on UWS patients with a confirmed PET negative (PET-) diagnosis (i.e., a complete bilateral hypometabolism of the associative frontoparietal cortex) on the one hand and either healthy controls or MCS+ patients on the other. The MCS+ patients here function as a conscious control group who still suffer from brain damage. We then test the models on datasets from “covertly aware” MCS* patients (partially preserved metabolism and activity within these areas). We hypothesize that if our modelled effects are valid, and if the sustained PET metabolism reflects covert awareness in the MCS* patients, our model should classify these patients as healthy controls/MCS+ rather than UWS PET-.

## 2 Results

### 2.1 Dynamic causal modeling and parametric empirical Bayes

Our first goal was to investigate the effective connectivity modulations best explaining the difference between healthy controls, UWS PET-, and MCS+ patients. We modelled time-series recorded from the three groups with DCM for CSD at a single-subject level, followed by PEB at the group-level. In doing so, we estimated the change in effective connectivity in 12 inter-node connections in the DMN, contrasting 11 healthy controls both with 6 UWS PET–patients and with 12 MCS+ patients, and the 12 MCS+ patients with 6 UWS PET-patients.

Following the inversion of the between-groups PEB model, a greedy search was implemented to prune away connections not contributing significantly to the free energy using BMR. Figure 4 shows the most parsimonious models and figure 5 shows the estimated log scaling parameters contrasting healthy controls with UWS PET-(A), MCS+ with UWS PET-(B), and finally, healthy controls with MCS+ (C). Here, we applied a threshold of >.99 for the posterior probability; in other words, connections that were pruned by BMR and connections with lower than .99 posterior probability with their respective log scaling parameter are faded out (figures 5A, 5B, 5C).

**Figure 1.**
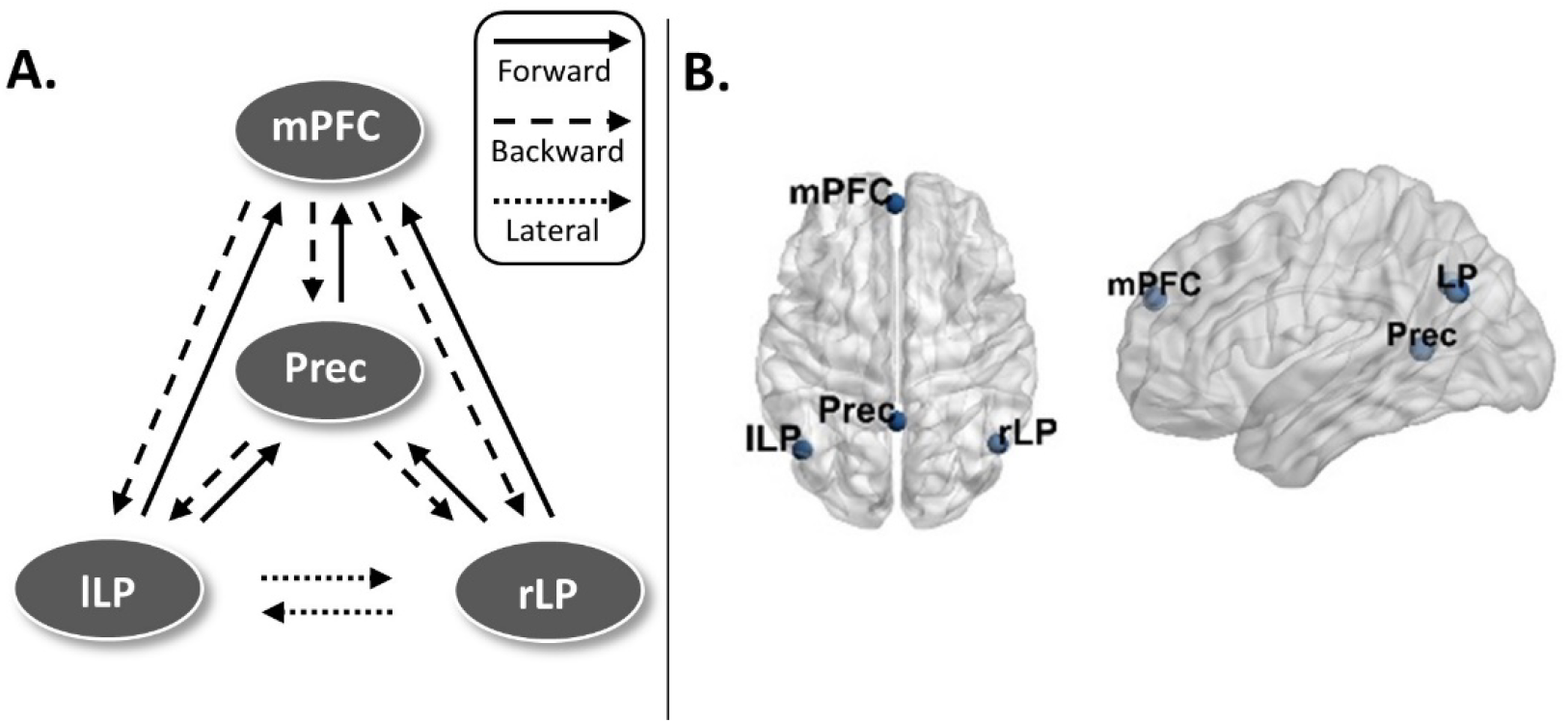
A. The fully connected, schematic representation of the default mode network (DMN). **B.** The node locations for the DMN. mPFC – medial prefrontal cortex, Prec – posterior cingulate cortex/precuneus, lLP – left lateral parietal cortex, rLP – right lateral parietal cortex.

**Figure 2.**
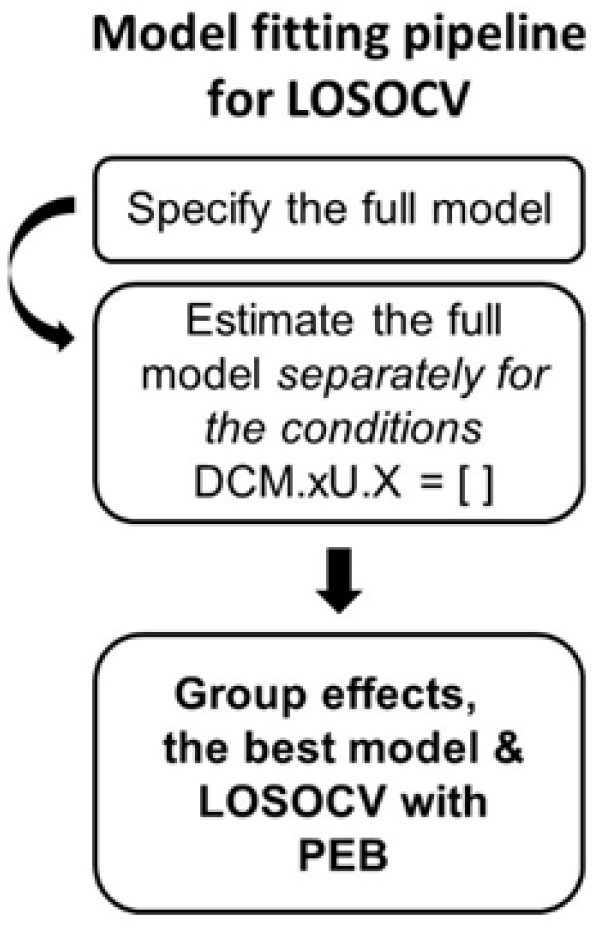
The pipeline for inverting the dynamic causal modelling (DCM) model for different subject-groups. This was done to find the best models for each patient group, to estimate th effective connectivity modulations between the patient groups, and as a prerequisite for th leave-one-subject-out cross-validation (LOSOCV) classification with parametric empirical Bayes (PEB) modelling.

**Figure 3.**
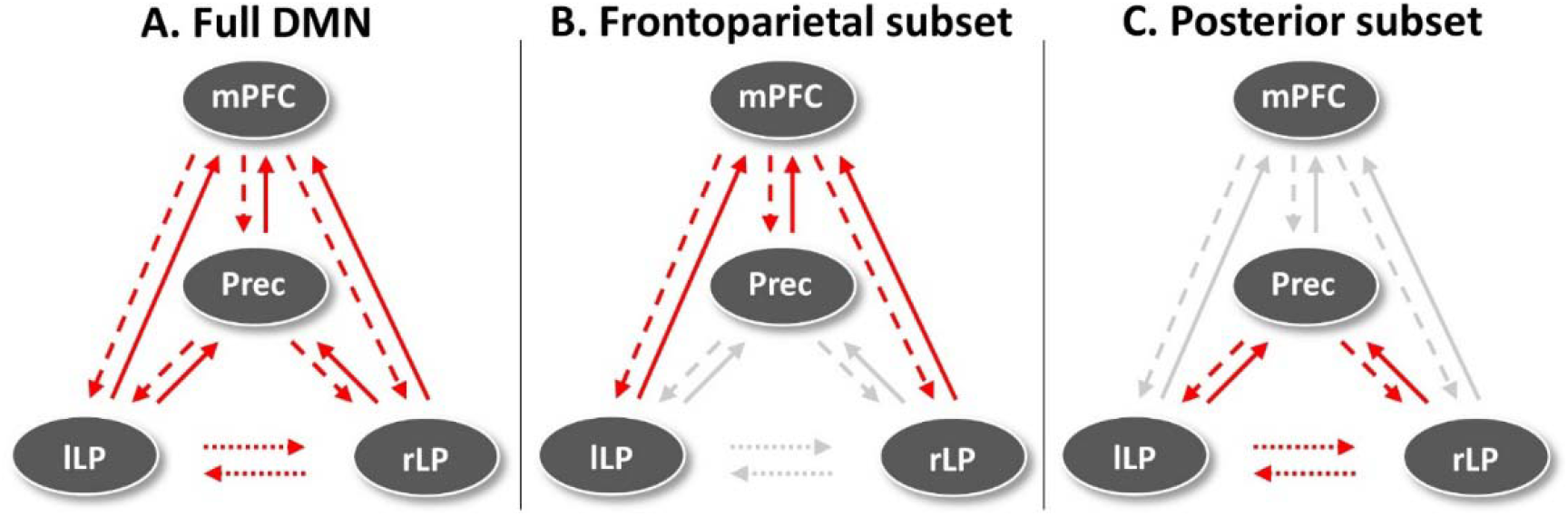
The hypothesis-driven subsets for the LOSOCV-paradigm. The red arrows indicate the connections included in each subset, and the grey arrows the left-out connections. First, predictions based on all connections were estimated (A). Next, predictions based on two connection subsets – frontoparietal (B) and parietal subsets (C) – were estimated. Lastly, we estimated predictions based on single connections in a data-driven approach. mPFC – medial prefrontal cortex, Prec – posterior cingulate cortex/precuneus, lLP – left lateral parietal cortex, rLP – right lateral parietal cortex, DMN – default mode network.

**Figure 4.**
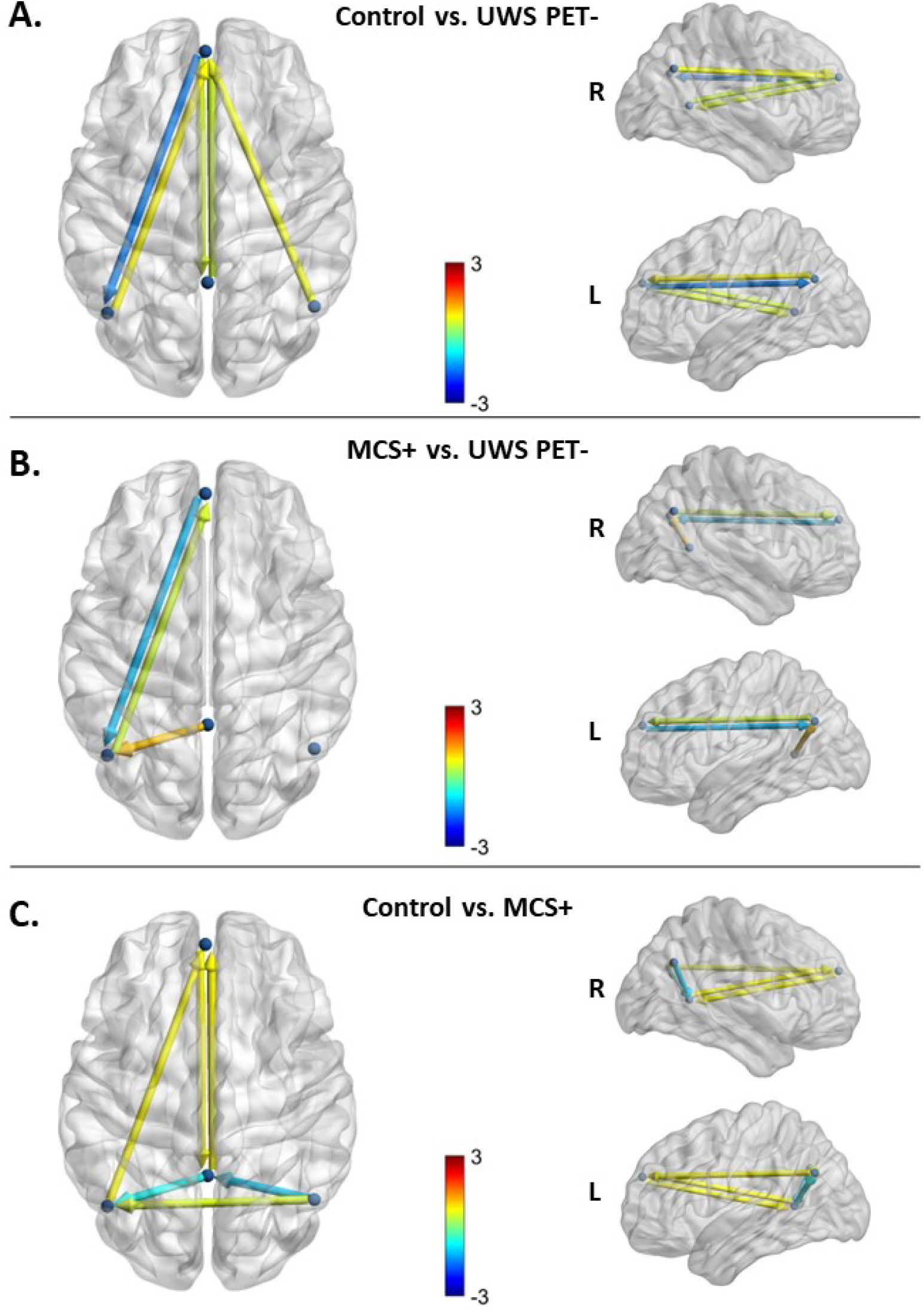
The most parsimonious DMN models after BMA and BMR contrasting the healthy controls (HC) and the UWS PET-, MCS+ patients and UWS PET-, and healthy controls and MCS+. Color shows modulation strength and direction. All panels express the modulations of couplings for the latter state relative to the first. **A.** The most parsimonious model best explaining the difference between healthy controls and UWS PET-patients. Three connections were pruned with an additional four having lower than .99 posterior probability of being present. All but one pruned connection was located between lateral parietal and PCC/precuneus nodes. When moving from the state of healthy controls to UWS PET-, the largest reduction in effective connectivity was in the backward connection from the medial prefrontal cortex to left lateral parietal cortex. **B.** The most parsimonious model best explaining the difference between the MCS+ and UWS PET-patients. Six connections were pruned by the BMR with an additional three connections having lower than .99 posterior probability of being present. When moving from UWS PET– to MCS+, the largest reduction was observed on the backward connection from the medial prefrontal cortex to left lateral parietal cortex, similar to the reduction between healthy controls and UWS PET^-^ patients. **C.** The most parsimonious model explaining the difference between healthy controls and MCS+ patients. Two connections were pruned with additional four connections having lower than .99 posterior probability of being present. When moving from the state of healthy controls to MCS+, the largest reductions on effective connectivity were on posterior connections between the lateral parietal cortices and PCC/precuneus.

**Figure 5.**
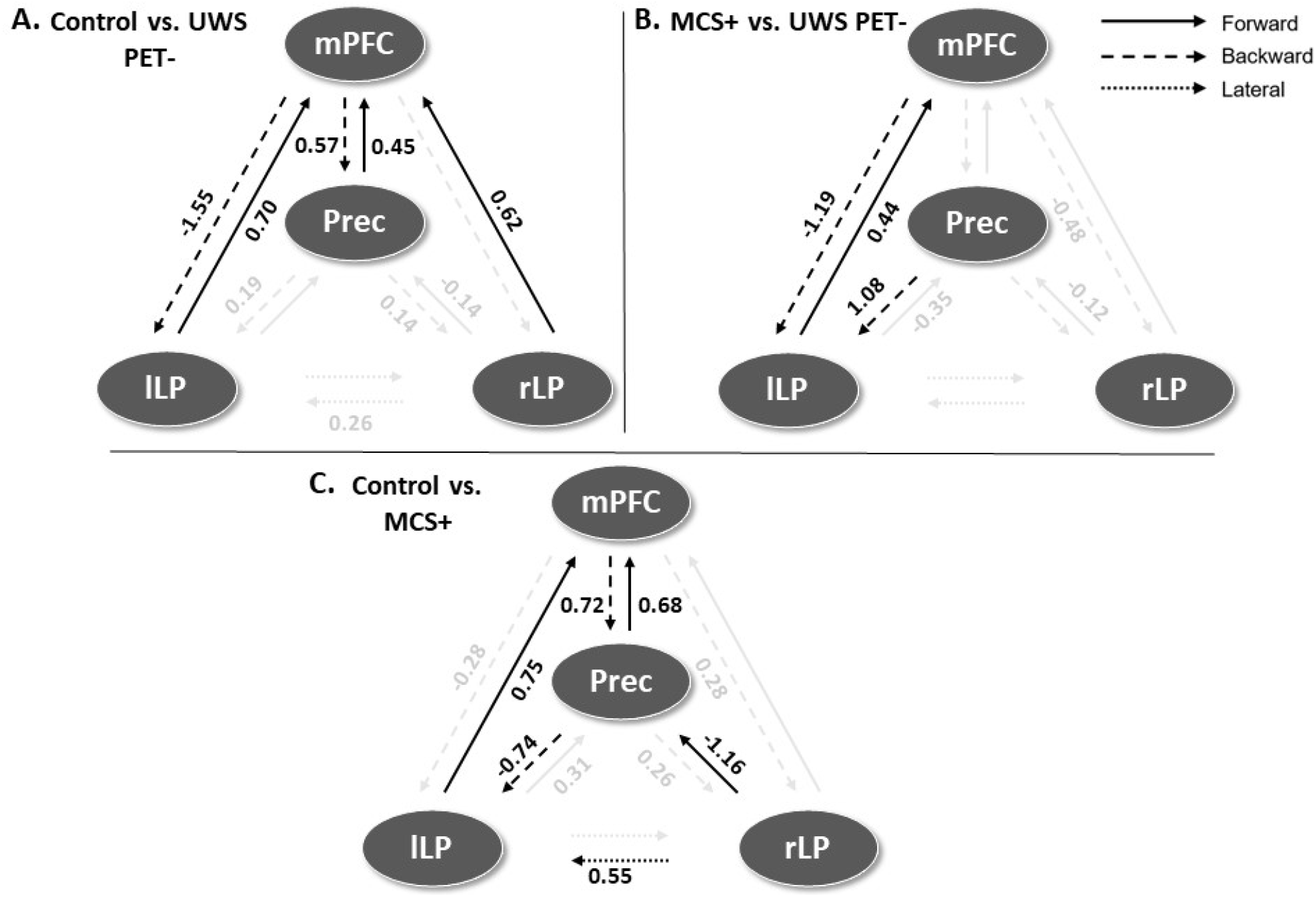
The log scaling parameters for the connection strengths in the DMN after BMR and BMA. Positive values represent an increase and negative values a decrease in effective connectivity for the three group comparisons. Connections that were pruned by BMR and connections with lower than .99 posterior probability with their respective log scaling parameter are faded out. **A.** The modulatory effects best explaining the difference between healthy controls (HC) and UWS PET– patients. Connectivity between lateral parietal and PCC/precuneus nodes were either pruned away by BMR or had lower than .99 posterior probability with low modulatory effects. The largest reduction was found on backward lateral frontoparietal connection from medial prefrontal cortex to left lateral parietal cortex. **B.** The modulatory effects best explaining the difference between MCS+ and UWS PET-. The modulatory effects on left bidirectional frontoparietal connections were both in the same direction as when comparing healthy controls to UWS PET-, with the largest reduction on left backward frontoparietal connection. In addition, right backward frontoparietal connectivity and posterior forward connectivity between lateral parietal nodes and PCC/precuneus reduced (smaller effect sizes), albeit with lower than .99 posterior probability of being present. **C.** The modulatory effects best explaining the difference between healthy controls and MCS+. The largest reductions were between the posterior nodes, between the lateral parietal nodes and PCC/precuneus. Bidirectional medial frontoparietal connectivity was increased in MCS+ in comparison to healthy controls. mPFC – medial prefrontal cortex, Prec – posterior cingulate cortex/precuneus, lLP – left lateral parietal cortex, rLP – right lateral parietal cortex.

On inverting the DMN for the control and UWS PET– groups, 3 connections were pruned away by BMR with additional 4 connections having lower than .99 posterior probability (figures 4A and 5A). All but one of the pruned connections were located within the posterior cortices between lateral parietal cortices and PCC/precuneus (except for the right backward frontoparietal connection). The largest reduction in effective connectivity was located on left frontoparietal connection; the backward connection between mPFC and left lateral parietal node was largely diminished for the UWS PET– group in comparison to healthy controls.

On inverting the DMN contrasting MCS+ and UWS PET-, only three connections survived the BMR process with at least .99 posterior probability (with additional three connections surviving pruning with lower than .99 posterior probability; figures 4B and 5B). As with the control vs. UWS PET– contrast, the largest reduction was on the left backward connectivity from mPFC to lLP, with left lLP-mPFC forward connectivity increasing.

On inverting the DMN for the contrast between healthy controls and MCS+, two connections were pruned by the BMR with additional 4 connections having lower than .99 posterior probability for being present (figures 4C and 5C). The largest reductions were between the posterior nodes, to and from the lateral parietal cortices and PCC/precuneus. In addition, the left frontoparietal backward connectivity was reduced, although with smaller than .99 posterior probability and with clearly smaller effect size than with UWS PET-. Other non-pruned connections were associated with small to medium increases.

In addition, we also observed increased connectivity (relatively small effect sizes) in most of the other connections with at least .99 posterior probability.

### 2.2 Leave-one-subject-out cross-validation

To conduct LOSOCV, the DCM model was inverted again, this time separately for each patient group. Following the inversion process, PEB was conducted repeatedly on the training set in each cross-validation run alongside LOSOCV analysis to generate the posterior probabilities for group-membership (see Methods).

First, the UWS PET– patients were classified alongside the controls based on the full DMN model, and two hypothesis-driven connection subsets (frontoparietal– and parietal connections; figure 6). A similar approach was applied classifying MCS+ patients alongside UWS PET– patients, and finally, healthy controls alongside MCS+ patients. Figures 6 and 7 show violin plots representing the individual posterior probabilities for the hypothesis-driven classifications and data-driven approach, for all three contrasts, respectively (A: control vs. UWS PET-, B: MCS+ vs. UWS PET-, C: control vs. MCS+). As seen in figure 6, frontoparietal connections classified correctly most of the controls and MCS+, and around half of the UWS PET-patients in the controls vs. UWS PET– and MCS+ vs. UWS PET-contrasts. Both full DMN and parietal subsets clustered most of the subjects around the chance level of 0.5. We further produced confusion matrices of prediction accuracy calculated by labelling posterior probabilities greater than 0.5 as a positive classification (for both hypothesis-driven subsets and the data-driven approach). See Supplementary Materials and figures s3 and s4.

**Figure 6.**
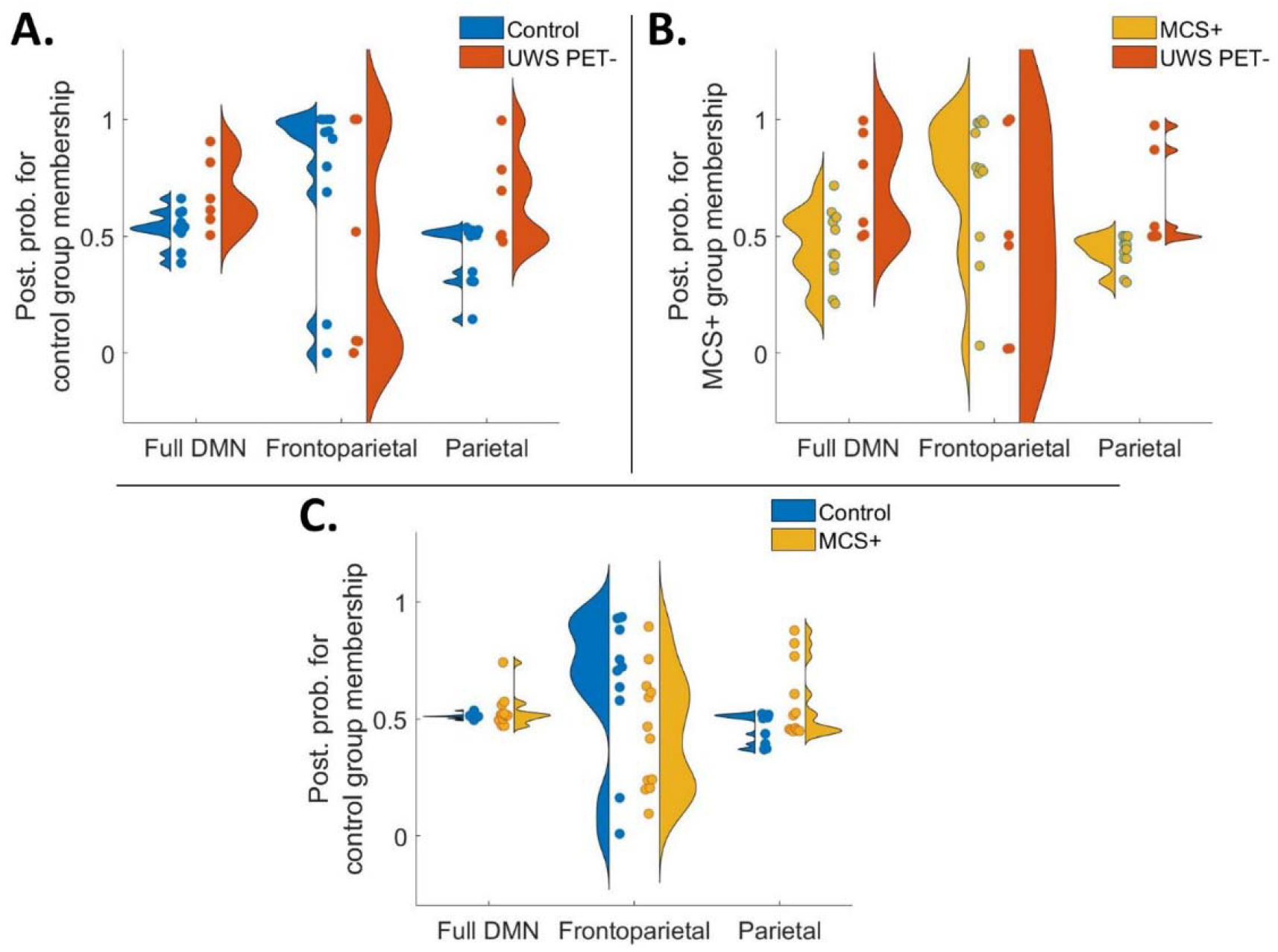
Violin plots representing diversity in posterior probabilities for healthy control group membership (A and C) and for MCS+ group membership (B) for the hypothesis-driven subsets for all three group contrasts. Each colored point specifies a subject. In a perfect model in panels A and B, the UWS PET-patients (*N* = 6), and in panel C, the MCS+ patients (*N* = 12) should approach a posterior probability of zero. Overall, the results show a trend for frontoparietal connections producing the best predictions. **A.** When classifying UWS PET-patients alongside healthy controls, the frontoparietal subset produced the best results. The individual data points reveal more consistent classifications of healthy controls. On all three panels, full DMN model and parietal subset produced classifications with most posterior probabilities bordering the 0.5 chance level. **B.** As in panel A, the best predictions when classifying UWS PET-patients alongside MCS+ were based on frontoparietal connections, specifically with MCS+ patients. **C.** Classification of MCS+ alongside healthy controls. Frontoparietal subset produced the best predictions, however with large variability on the performance across the subjects.

**Figure 7.**
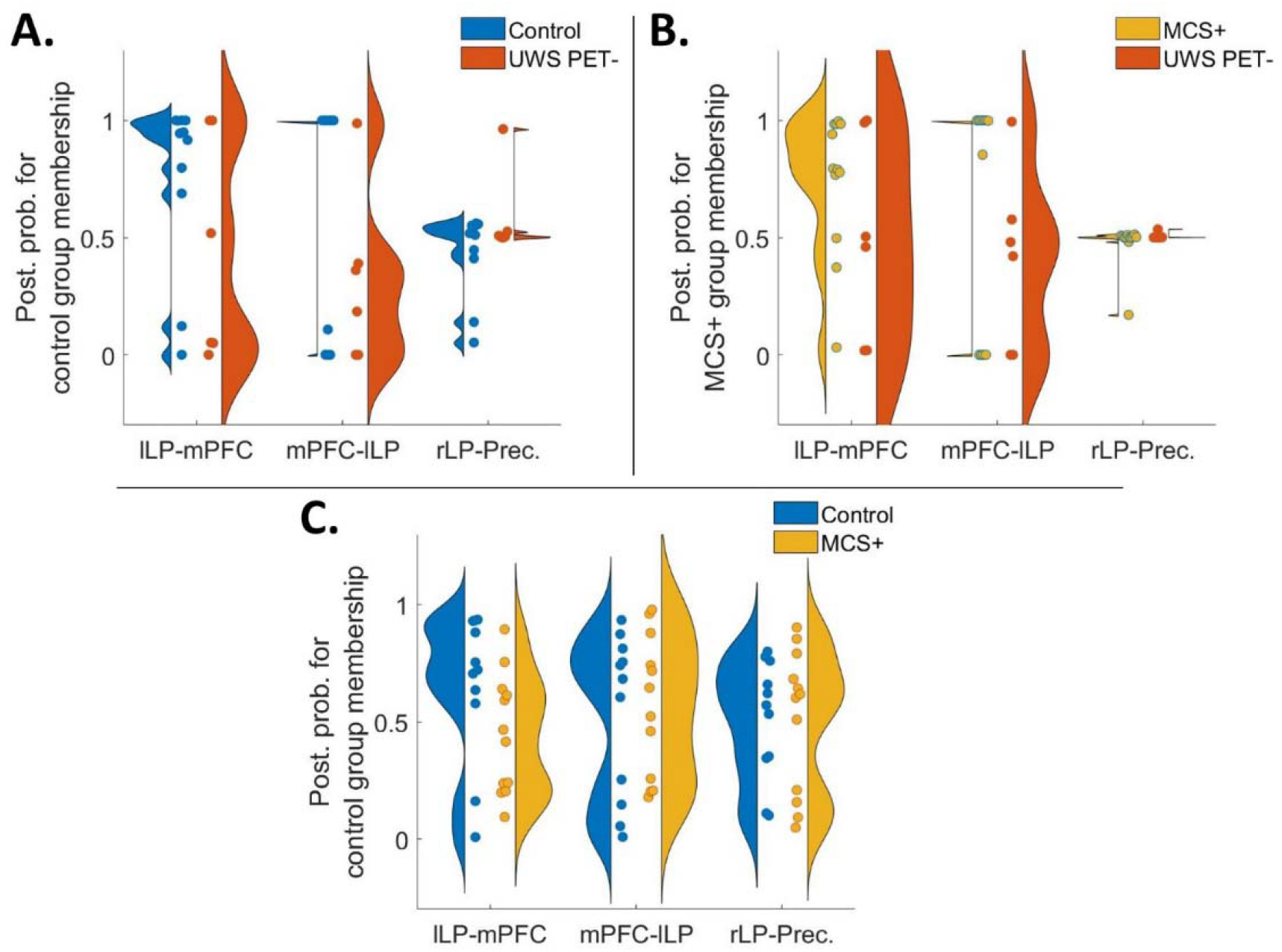
Violin plots representing diversity in posterior probabilities for control group membership (A and C) and for MCS+ group membership (B) for the data-driven connections for all three contrasts. Each colored point specifies a subject. In a perfect model in panels A and B, the UWS PET-patients, and in panel C, the MCS+ patients should approach a posterior probability of zero. Overall, the best predictive performance was based on the left bi-directional frontoparietal connections when classifying UWS PET-alongside controls (A) and MCS+ (B). Largest inconsistencies and variability were on classifications of MCS+ alongside healthy controls. **A.** Left frontoparietal connectivity from mPFC to lLP produced the best predictions (mean posterior probabilities) of the group-membership when classifying UWS PET-alongside healthy controls. As with the hypothesis-driven subsets, the classifications were more accurate with healthy controls than with patients. **B.** As in panel A with healthy controls and UWS PET-patients, the classification performances based on mPFC-lLP and lLP-mPFC produced the most consistent results when contrasting UWS PET-patients alongside MCS+. **C.** Mean posterior probabilities for classification of MCS+ alongside healthy controls. The performance of the models based on the single connections did not produce consistently accurate classifications. mPFC – medial prefrontal cortex, Prec – posterior cingulate cortex/precuneus, lLP – left lateral parietal cortex, rLP – right lateral parietal cortex.

We then moved to a data-driven approach in which we first predicted the patient group membership based on the connections with the largest reductions in PEB, one at a time, working through all connections. Lastly, we checked combinations based on the connections’ respective classification accuracies (see Methods). The bi-directional left frontoparietal connectivity produced the best predictions, especially when classifying the UWS PET-from both, healthy controls and the MCS+ patients (figures 7A and 7B), with the best predictions based on the backward mPFC-lLP connectivity. Forward lLP-mPFC connectivity produced the best predictions for controls vs. MCS+ contrast, especially for the healthy controls (7C). None of the tested combinations improved classification performance.

### 2.3 Leave-one-state-out cross-validation

Finally, the predictive power of DCM modelling was generalized in two more difficult classification problems; each model was trained first on healthy controls and UWS PET– and then tested on the previously unseen UWS PET + group. A similar approach was used with a training set consisting of MCS+ and UWS PET-patients. The individual posterior probabilities for the five UWS PET + patients represented in a violin plot for both, the hypothesis subsets (panels A and B; controls vs. UWS PET– and controls vs. MCS+, respectively) and for data-driven connections (panels C and D) are shown in figure 8. The hypothesis-driven subsets did not classify the MCS* as controls or MCS+. Instead, when trained on datasets from healthy controls and UWS PET-, the frontoparietal subset classified four out of five MCS* patients as UWS PET-.

**Figure 8.**
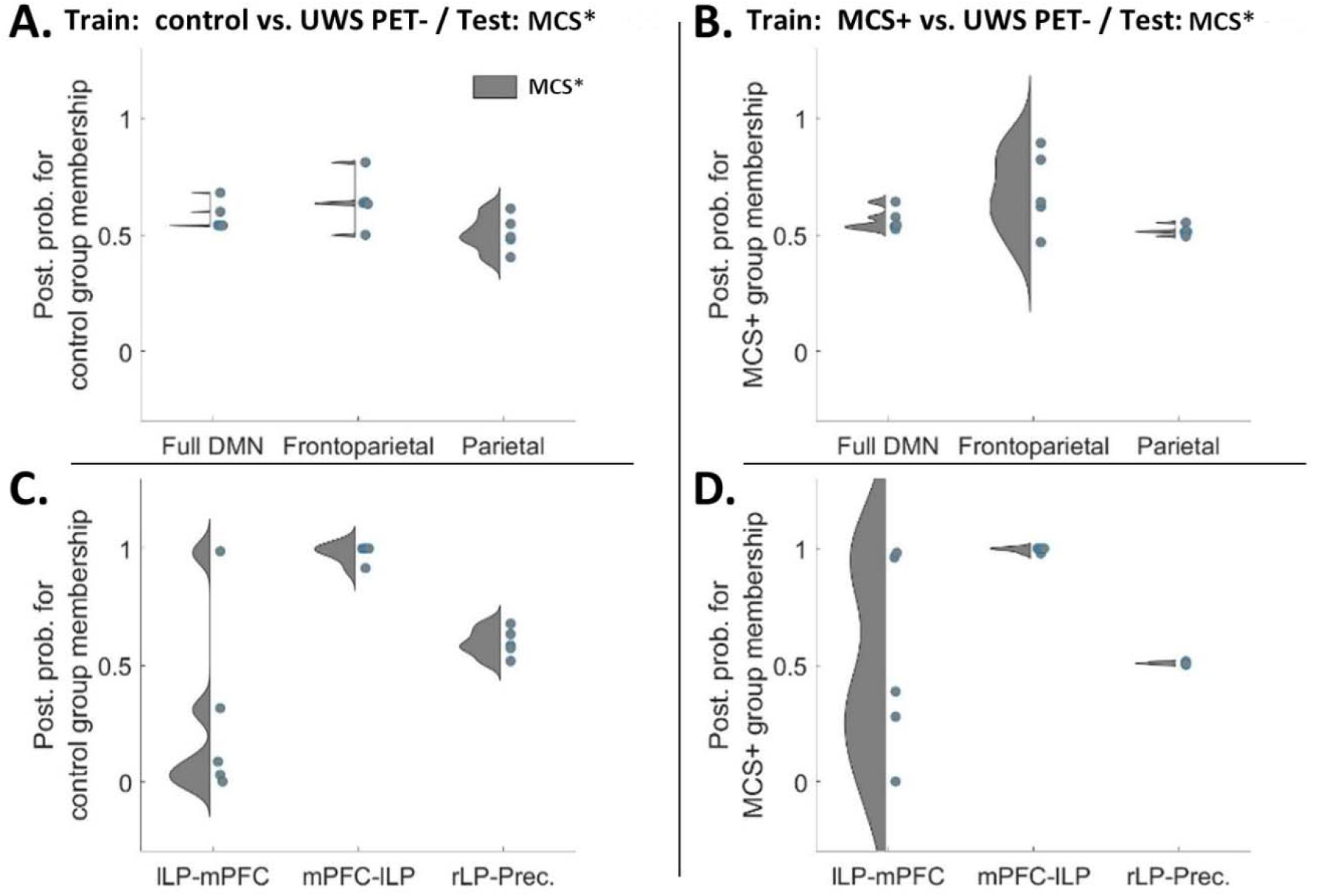
Violin plots representing diversity in posterior probabilities for control group membership (A and C) and for MCS+ group membership (B and D) for both, the hypothesis– and data-driven predictions. Here, the models were trained on datasets from controls and UWS PET-(A & C) or from MCS+ and UWS PET-(B and D) and tested on unseen data from MCS* patients (*N* = 5). Each colored point specifies a subject. Overall, the left backward frontoparietal connectivity produced the best group membership predictions. **A and B.** Mean posterior probabilities for classification of MCS* patients alongside healthy controls (A) and MCS+ patients (B). Neither of the hypothesis-driven subsets, nor the full DMN, clearly classified the unseen MCS* patients as members of either train-group. **C and D.** Left frontoparietal connectivity from mPFC to lLP produced almost perfect predictions for the MCS*, classifying all five patients as either controls or MCS+ rather than UWS PET-patients. Unlike the backward connectivity, predictions based on the left forward connectivity from lLP to mPFC, the model classified four of five MCS* patients as UWS PET-rather than as controls (C). Similarly, when trained on MCS+ and UWS PET-, the model classified three of five patients as UWS PET-rather than MCS+. mPFC – medial prefrontal cortex, Prec – posterior cingulate cortex/precuneus, lLP – left lateral parietal cortex, rLP – right lateral parietal cortex.

With the data-driven approach, the left backward frontoparietal connectivity from mPFC to lLP produced nearly perfect predictions classifying the MCS* datasets as controls (8C, p(control|MCS* > .92) and MCS+ (8D, p(MCS+|MCS* > .98) rather than as UWS PET-group. Similar as with frontoparietal subset (8A), the left forward connectivity from lLP to mPFC classified four of five patients as UWS PET-patients rather than healthy controls – three of them with a high posterior probability. This dissociation was not as prominent when training the model with MCS+ and UWS PET-patients (8D); the backward connectivity still produced nearly perfect classifications (of MCS* as MSC+) while the predictions based on the forward connectivity showed larger variability.

## 3 Discussion

In this cross-sectional, retrospective analysis, we applied spectral DCM to high-density EEG data with PEB to investigate the difference in effective connectivity dynamics between cortico-cortical regions of the DMN in DoC patients (UWS, MCS+, and MCS*) and healthy controls. Overall, the modelling results indicate a key difference between healthy controls or MCS+ patients and unresponsive patients with compatible hypometabolism (UWS PET-) in left-hemispheric backward frontoparietal connectivity. Furthermore, with out-of-sample cross-validation, we demonstrated that this association is robust enough to not only distinguish patient groups from each other, but also generalizes to an unseen data subset, collected from seemingly unresponsive patients showing preserved brain activity compatible with MCS (MCS*). These results identify specific alterations in the DMN after severe brain injury and highlight the clinical utility of EEG-based measurement of effective connectivity for identifying covert consciousness.

### 3.1 Dynamic causal modelling

The most parsimonious model explaining the difference between healthy controls and unresponsive patients with congruent hypometabolism (UWS PET-) indicated a large relative reduction in left-hemisphere backward frontoparietal connectivity in UWS PET-patients. Additionally, a small, lower-probability reduction from right lateral parietal cortex to precuneus was found. Interestingly, excluding the right parietal connection, the estimated connectivity in the posterior nodes – within the ‘posterior hot zone of conscious contents’ (Koch et al., 2016; Siclari et al., 2017) – was either pruned away from the model best explaining the difference or returned only small, low-probability increases suggesting lower relative importance for the posterior hot zone in explaining the difference between healthy controls and UWS PET-patients. Generally, connections pruned by BMR are considered not to contribute towards the model evidence (Zeidman et al., 2019). In contrast, in a previous fMRI DCM study with DoC patients, precuneus/PCC-related connectivity reduction was the key difference; specifically, the recurrent connectivity (down-regulation of the PCC itself) was found to be diminished on UWS patients in comparison to both, MCS patients and healthy controls (Crone et al., 2015).

However, not only are the data between Crone et al.’s (2015) study and the present study from different modalities, and thus, direct comparisons of the results unsound, the underlying neurobiologically motivated models used by DCM are different for hemodynamic vs. electrophysiological data leading to a different interpretation of the modulatory effects. The interpretation of the modulatory effects in DCM for fMRI vs. EEG differ in that positive and negative values indicate excitatory and inhibitory effects in fMRI data (except for recurrent connections, for which the connection strength is always negative and modulations reflect increases or decreases in comparison to the prior). In DCM for EEG, positive modulations indicate an increase and negative a decrease in connectivity relative to the prior. In the neural mass model we used here, the backward connections are thought to have more inhibitory and largely modulatory effect in the nodes they target (top-down connections), while forward connections are viewed as having a strong driving effect (bottom-up; Salin & Bullier, 1995; Sherman & Guillery, 1998).

Here, we modelled the data with the default ERP neuronal model (David et al., 2005) primarily in order to produce comparable results with prior DCM for EEG work modelling consciousness (Boly et al., 2011a; Boly et al., 2012; Ihalainen et al., 2021). Further, we aimed to model disrupted consciousness at the level of active networks rather than focusing on e.g., synaptic hypotheses or recurrent self-connections (intrinsic connectivity), which could be better captured with other neuronal models such as the local field potential model (Moran et al., 2007) or the Canonical Microcircuits model (Bastos et al., 2012; see also Moran et al., 2013 for a review of neuronal population models). Hence, we only estimated extrinsic connectivity – i.e., connectivity between cortical areas. It is possible that the observed differences in the network dynamics are driven by modulations in self-inhibiting, recurrent connectivity within the cortical sources or within and between subcortical networks driving the disruptions in the DMN (Chen et al., 2018; Coulborn et al., 2021). A worthwhile endeavor for future DCM for EEG studies would be to model the extent to which recurrent, within-source cortical connectivity may or may not drive the modulations in extrinsic connectivity.

To the best of our knowledge, only one study has used DCM for EEG in DoC populations. Boly and colleagues (2011a) showed that in an auditory mismatch negativity paradigm, the difference between UWS, MCS, and healthy controls was an impairment of backward connectivity from frontal to temporal cortices in the UWS patients, emphasizing the importance of top-down processing for conscious perception. Similarly, in the present resting-paradigm, the key difference distinguishing UWS PET-from both, MCS+ and healthy controls was decreased left-lateralized backward frontoparietal connectivity in UWS PET-patients, although from medial prefrontal cortex to lateral parietal cortex and not to superior parietal cortex. It is important to note, however, that the differences in the paradigm, methodology and in the models estimated render rigorous, direct comparisons of the results between Boly et al.’s (2011a) study and the present study infeasible. Moreover, even though some of the patients may have been the same between the present study and those of Boly et al. (2011a), the data were different (here recorded with high-density EEG during PET while Boly et al. (2011a) used a 64 channel EEG prior to TMS-EEG). Despite the methodological differences, the results of both studies suggest a crucial role for lateral backward connectivity originating from the frontal cortex. Future studies should investigate this further by modelling the connectivity related to temporal areas, and the backward frontoposterior connectivity in other resting networks (see below) in DoC patients.

Like with healthy controls vs. UWS PET-comparison, the largest difference between MCS+ and UWS PET-patients was left frontoparietal backward connectivity, with UWS PET-patients again showing reduced connectivity. Furthermore, the left forward parietofrontal connectivity and backward connectivity from precuneus to lLP both increased, reproducing the modulations between healthy controls and UWS PET-. These changes were accompanied by smaller, low-probability reductions in forward connectivity within the posterior hot zone and in right backward frontoparietal connectivity. These modelling results highlight the left frontoparietal backward connectivity as the key distinguishing difference when comparing healthy controls or conscious patients with unconscious patients and complement those of previous studies discriminating DoC patients from scalp-level EEG connectivity; especially frontal and parietal functional connectivity has been shown to consistently discriminate DoC patients (Chennu et al., 2014; Chennu et al., 2017). As the direction and the spatial location of the changes in connectivity were similar with the comparisons involving UWS PET-patients, and distinguishable from those when comparing healthy controls vs. MCS+, we were motivated to further test the predictive power of the modulatory effects (see below).

In contrast, the largest connectivity reductions between healthy controls and MCS+, although relatively smaller than in previous contrasts, were associated with the precuneus node in the posterior hot zone. The left backward frontoparietal connectivity was again reduced, but by a smaller effect and with lower than .99 probability of being present in the most parsimonious model. The activity changes in the posterior hot zone of conscious content have been associated with changes in consciousness not only during sleep (Lee et al., 2019; Siclari et al., 2017) and general anesthesia (Alkire et al., 2008; Ihalainen et al., 2021), but also in DoC patients (Vanhaudenhuyse et al., 2010; Wu et al., 2015). Moreover, previous studies have suggested a subdivision of the frontoparietal network into two anticorrelated subnetworks; an “intrinsic” network encompassing precuneus/PCC, anterior cingulate/mesofrontal cortices, and parahippocampal areas associated with internal awareness, and into an “extrinsic” central executive network encompassing dorsolateral prefrontal and posterior parietal areas linked with the intensity of external awareness (Boveroux et al., 2010; Vanhaudenhuyse et al., 2011). The observed decrease in the left lateral frontoparietal connectivity in the present study between UWS PET-patients and healthy controls or MCS+ patients may reflect such diminished internal awareness in the UWS PET-patients. To that end, future endeavors should investigate the modulatory effects and the possible predictive power of such modulations in other resting state networks, such as the central executive network.

However, changes in the physiological state of the frontoparietal network alter not only consciousness but also several other brain functions such as vigilance and attention (Hohwy, 2009; Koch et al., 2016). Moreover, the specific areas of the DMN have been associated with specific cognitive functions; for example, the frontal areas seem to be important for self-reference, whereas the precuneus/PCC in autobiographical memory (Whitfield-Gabrieli et al., 2011). It remains a possibility that the found modulations in the DMN reflect changes also in other cognitive functions, rather than in awareness alone.

It is important to bear in mind that DoC patients typically suffer from widespread structural brain damage often accompanied by distributed white matter anomalies (Annen et al., 2018; Fernández-Espejo et al., 2012; Tshibanda et al., 2009). Hence, it is relevant to consider the feasibility and validity of applying DCM to DoC patients; this is particularly true for DCM for EEG which requires the specification of the anatomical locations of the nodes/sources a priori, and with patients with non-traumatic etiology, e.g., patients with anoxic brain damage (King et al., 2011; Boly et al., 2011b). Here, we mitigated these concerns first by limiting our modelling to DMN, a resting state network previously associated with DoC (Boly et al., 2008, 2009; Crone et al., 2011; Crone et al., 2015; Fernández-Espejo et al., 2012; Heine et al., 2012; Laureys et al., 1999; Laureys, 2005; Vanhaudenhuyse et al., 2010) and with changes in the conscious state e.g., due to anesthesia (Boveroux et al., 2010; Greicius et al., 2008; Stamatakis et al., 2010) and sleep (Horovitz et al., 2009).

Second, we selected only patients with TBI as compared to non-traumatic aetiologias, it has been associated with more focal injury centered often on areas susceptible to rotational forces, such as the brainstem, midbrain, thalamus, hypothalamus, cerebellum, and posterior corpus callosum (Guldenmund et al., 2016; Newcombe et al., 2010). That said, future studies should look to extend these results to other aetiologias; an obvious downside for trying to control for the individual differences due to brain damage by restricting the analysis to TBI patients only, was reduced sample size. By including other aetiologias, future studies could not only aim to replicate and verify these results with larger samples and better power, but to increase the potential clinical utility by extending them to cover larger patient populations.

Third, we applied a special case of Bayesian model selection (BMS), Bayesian model reduction (BMR), to invert multiple nested models from a single, fully connected DMN (see Methods). A particular advantage here is that BMR can be applied using an explanatory approach, in which no strong a priori hypotheses about the model parameters are needed. This enables a greedy search to compare the negative free energies of the reduced (nested) models by iteratively discarding parameters that do not contribute to the free energy. The procedure stops when discarding any parameters starts to decrease the negative free energy, returning the model that most effectively trades-off goodness of fit and model complexity in explaining the data. BMR applied in this way allows one to estimate a large model space from a single, specified full model in a relatively short period of time (Friston & Penny, 2011; Rosa et al., 2012; Zeidman et al., 2019).

Nevertheless, it is possible that not all true influences on the specific regions are captured by the specified full model. Moreover, the explanatory approach to BMR is conducted under the assumption that all reduced models are equally probable a priori, and thus, the full model should only contain parameters that are biologically plausible. Here, we cannot exclude physical damage to cortical areas and pathways crucial to the functioning of the DMN.

That said, our aim was not to make any strong claims about the “true” model; to draw stronger conclusions about the “true” underlying neuronal basis using DCM for EEG, structural MRI imaging assessing the extent of the damage in specific patients, possibly in adjunct with source localization of the EEG signals, should be applied. Here, the aim was to demonstrate and to compare the predictive performance of effective connectivity in the clinical context. Additionally, demonstrating predictive value with significant generalization performance with cross-validation, the level of confidence we can ascribe to our modelling results increases.

### 3.2 Leave-one-subject/state-out cross-validation (LOSOCV)

To test whether the effective connectivity modulations were consistent enough across the patient groups and healthy controls to reliably distinguish the groups from each other, we first conducted a leave-one-subject-out cross-validation based on hypothesis-(full DMN, frontoparietal, and parietal subsets; figure 6) and data-driven subsets of connections (figure 7). Amongst the connection subsets, the frontoparietal connections performed the best, classifying most controls and MCS+, and half of the MCS* patients correctly in the healthy controls vs. UWS PET– and in MCS+ vs. UWS PET-contrasts. The full DMN and parietal subsets clustered most of the subjects around the chance level in all three contrasts.

We then moved to a data-driven approach in which we first predicted the patient group membership based on the connection with the largest reduction in PEB, one at a time, working through all connections. It is important to note that searching for the best connection in this way increases the risk of overfitting the model by potentially extracting some of the residual variation – noise – as if representing the underlying model structure. However, the fact that the best model generalized to an unseen dataset suggests that the results may reflect a genuine effect (see below).

With the data-driven approach, the bi-directional left frontoparietal connectivity produced the best predictions, especially when classifying the UWS PET-from both, healthy controls and the MCS+ patients. The single best performing connection was the backward frontoparietal connectivity (figure 7). Not surprisingly, the classifications were more accurate and consistent with healthy controls than with patients; classifications of patients suffering from severe brain damage, and thus, from highly disrupted brain functioning, were expected to vary more than those of healthy controls. Next, we combined the single connections into data-driven subsets based on the classification performance: none of the combinations improved the performance of the single connections.

Last, the predictive power of DCM modelling was generalized in two more difficult classification problems; following the hypothesis– and data-driven approaches above, we trained each model on healthy controls or MCS+ on the one hand, and UWS PET-patients on the other, and then tested the models on the previously unseen, “covertly aware” MCS* patients. The hypothesis-driven subsets did not classify the MCS* patients as controls or MCS+. Crucially, with the data-driven approach, the left backward frontoparietal connection produced nearly perfect predictions classifying all five patients as either controls or MCS+ rather than UWS PET-patients. These results are compatible with previous PET imaging results by Thibaut and colleagues (2021) who observed higher brain metabolism in the lateral and medial frontoparietal network in MCS* patients when compared to UWS PET-patients. Interestingly, their resting state EEG results with functional connectivity indicated a difference in the left hemisphere (and at the whole brain level) when comparing MCS* to both UWS PET– and to MCS patients. To that end, alpha connectivity was higher and delta connectivity lower in MCS* when compared to UWS PET-patients. Moreover, a difference between MCS* and MCS was observed in the left hemisphere with the latter having higher connectivity in the theta band. This finding further extended to MCS* vs. MCS+ comparison.

While the results of the cross-validation here should be interpreted with caution due to the relatively low number of subjects in our study, the results, in conjunction with those of Thibaut and colleagues (2021) suggest a pivotal role for left hemisphere connectivity in distinguishing MCS* from UWS PET– and MCS patients. Furthermore, the present results highlight the importance of the frontoparietal connectivity – particularly the left-lateralized backward connectivity – when predicting these states of consciousness. The importance of frontoparietal connections within the DMN for dissociating states of consciousness in DoC patients is not surprising given the previously established association between conscious awareness and the DMN (Boly et al., 2008, 2009; Boveroux et al., 2010; Vanhaudenhuyse et al., 2010). More specifically, consciousness is thought to require brain-wide broadcasting of information by a “global workspace” associated with brain areas within the frontoparietal network (Baars, 1988; Baars, 1997; Dehaene et al., 2011).

It is important to note, however, that the global neuronal workspace theory (GNW) is not a localisationist approach but rather posits a distributed “router” for conscious access (Dehaene et al., 2011). The extensive and rapid bidirectional connectivity between the hubs of the GNW is thought to trigger the sudden collective and coordinated activity mediating global broadcasting (Mashour et al., 2020). Aptly, these hubs initially included the prefrontal cortex and parietal cortex (in combination with a set of specialized and modular perceptual, motor, memory, evaluative, and attentional processors) although it has later been complemented with other, potentially equally important hubs (such as the anterior and posterior cingulate and the precuneus). The observation that the changes in the long-range frontoparietal connectivity best predicts the state of consciousness is in accordance with the suggested importance of the connectivity between the hubs in the GNW. This is in contrast with the more restricted, content-specific neural correlates of consciousness often associated with the posterior hot zone (Koch et al., 2016).

However, it is important to keep in mind that presumably, when the patient becomes “more” conscious, different content becomes more globally available for conscious processing throughout the brain, affecting and employing different cognitive systems (Hohwy, 2009). In other words, any major changes in the physiological state alter not only consciousness but other cognitive systems as well, many of which depend on levels of arousal-promoting neuromodulators. Therefore, it remains possible that the predictive performance of the frontoparietal effective connectivity is related not only to the state of consciousness, but also to other arousal-related cognitive processes. Further, it is worth noting that here, we limited our contrasts to MCS+, excluding minimally conscious – (negative) from the analyses. Our rationale was to include a second, irrefutably conscious control group. Nonetheless, future studies should include MCS-patients to better control for possible confounds of behavior and language functions present in MCS+.

In summary, our results indicate a key difference between healthy controls or MCS+ patients and unresponsive patients with congruent hypometabolism in left-lateralized backward frontoparietal connectivity. With out-of-sample cross-validation, we demonstrated that this association is robust enough to not only distinguish patient groups from each other, but also generalizes to an unseen data subset, collected from seemingly unresponsive patients. These results contribute towards identifying specific alterations in network interaction after severe brain injury, and importantly, suggest clinical utility of EEG-based effective connectivity in identifying covertly aware patients who seem behaviorally unresponsive.

## 4 Methods

### 4.1 Data Acquisition

We assessed effective connectivity within the DMN and whether modulation of this connectivity predicted states of consciousness in patients with DoC. The patients included were referred to the University and University Hospital of Liège (Coma Science Group and Centre du Cerveau²) from clinical centers across Europe since 2008. The data collection was approved by the Ethics Committee of the University Hospital of Liège and the patients’ legal guardians gave written informed consent. Data were also collected from healthy controls as a reference group, all of whom gave informed written consent before participation.

The dataset consisted of the patient data with 26 healthy controls (total *N* = 188). Parts of these data have already been published in previous studies (Carrière et al., 2020; Chennu et al., 2017; Panda et al., 2021). From the dataset, we identified patients admitted due to traumatic brain injury (TBI; *N* = 76). Amongst the TBI patients, we further identified those diagnosed with UWS (Laureys et al., 2010, *N* = 11) or MCS+ (Bruno et al., 2011, *N* = 12). Patients admitted due to any other etiology, e.g., anoxia or hemorrhage, and patients diagnosed with any other condition than UWS or MCS+, were excluded from the further analyses. See Supplementary materials for more details of the rationale and process of pruning the dataset. The patient groups were further divided based on their respective PET-scans – either into a PET-positive (PET+) or a PET-negative (PET-) sub-group (table 1). Amongst the healthy controls, using the random number generator in MATLAB, we (pseudo-randomly) drew a cohort of 11 control subjects to adjust for the group-size discrepancies. There were no meaningful differences in the mean ages between the groups (in a Bayesian ANOVA the probability for the model including the main effect of age: p(M|data) = 0.247, Bayes factor = 0.328).

**Table 1.** The mean age in years (SD), the total number of patients, and the number of PET+ and PET-of patients in each of the different DoC-groups. UWS – unresponsive wakefulness syndrome, MCS+ – minimally conscious state plus, TBI – traumatic brain injury.

| Patient group | Mean age (SD) in years | N<sup>TOTAL</sup> | PET+ | PET- | Etiology |
| --- | --- | --- | --- | --- | --- |
| UWS | 30.7 (8.5) | 11 | 5 | 6 | TBI |
| MCS+ | 38.3 (10.3) | 12 | 11 | 1 | TBI |
| Controls | 30.9 (6.7) | 11 | - | - | - |

Both PET scans and high-density EEG recordings were acquired at the same time and the patients were behaviorally assessed using the Coma Recovery Scale – Revised (CRS-R; Kalmar & Giacino, 2005) on the same day, before and after the scans (and on other days for a total of five assessments). Both patients and healthy volunteers were asked to stay awake during the data collection. The behaviorally apparent arousal levels of patients were monitored during the data collection session to ensure that they stayed awake and awaken with auditory/tactile stimulations if necessary.

PET scans were acquired and interpreted using methodology described in Stender et al. (2014) and in the Supplementary materials. Briefly, the analysis results were visually inspected by a trained clinician/researcher. Complete bilateral hypometabolism of the associative frontoparietal cortex without any voxels with preserved metabolism led to PET-diagnosis, whereas partial preservation with incomplete hypometabolism in these areas yielded a diagnosis of PET+ (Laureys et al., 2004; Nakayama et al., 2006; Thibaut et al., 2012).

The EEG data consisted of high-density EEG recordings of 20-30 minutes (256-channels, EGI), acquired during the F-fluorodeoxyglucose (FDG) uptake, just prior to the start of the PET imaging. The data were recorded at a sampling rate of either 250 Hz or 500 Hz (downsampled to 250 Hz). Data from the channels from the neck, cheeks, and forehead were discarded due to contributing most of the movement-related noise. We were left with the data from 173 channels on the scalp for further analysis. The raw signals were filtered from 0.5 – 45 Hz, with additional line noise removal at 50 Hz (notch-filter). We further restricted the DCM analysis to 1 – 30 Hz due to excessive high-frequency noise components. Via calculating the normalized variance, the excessively noisy channels and epochs were identified and either manually rejected or retained by visual inspection. Lastly, the data were re-referenced to a common average.

### 4.2 Dynamic causal modeling

We first imported the first 60 artefact-free 10-second epochs, in to SPM12 (Wellcome Trust Centre for Human Neuroimaging; www.fil.ion.ucl.ac.uk/spm/software/spm12). To analyze the resting effective connectivity within the DMN, DCM for EEG cross-spectral densities (CSD) was applied (Friston et al., 2012; Moran et al., 2009). Here, the observed cross-spectral densities in the resting-EEG are explained by a generative model that combines a biologically plausible neural mass model with an electrophysiological forward model mapping the underlying neural states to the observed data (ERP-model; Moran et al., 2013). The idea is to model the source activity over time in terms of causal relationships between interacting inhibitory and excitatory populations of neurons.

Each source – or node – is connected to each other via extrinsic connections, while each subpopulation within each source is connected to each other via intrinsic connections. Here, however, we aimed to model disrupted consciousness at the level of active networks, and hence, we estimated extrinsic connectvity between the nodes within the DMN. Among the extrinsic connectivity, the top-down – or backward – connections are thought to have inhibitory and modulatory effects on the nodes they target, while forward connections are viewed as having a strong excitatory driving effect (bottom-up; Salin & Bullier, 1995; Sherman & Guillery, 1998).

Within each node, second-order differential equations describe the hidden state of neural activity that depends on both the parameterized intrinsic and extrinsic connection strengths. This enables the computation of the linear mapping from the endogenous neuronal fluctuations to the EEG sensor spectral densities, and consequently, permits the modelling of differences in the spectra due to changes in the underlying neurophysiologically meaningful parameters. These parameters describe, for example, the intrinsic and extrinsic connectivity of coupled neuronal populations (i.e., sources) and their physiology. For further information about EEG DCM, see for example Friston et al. (2012), Kiebel et al. (2008), and Moran et al. (2009).

### 4.3 Model specification

Fitting an EEG DCM model requires the specification of the anatomical locations of the nodes/sources a priori. Here, we only model the DMN, which has been previously associated with DoC (Boly et al., 2008; Crone et al., 2011; Crone et al., 2015; Heine et al., 2012; Lin et al., 2017). The schematic representation and the node locations (adopted from Razi et al., 2017) are shown in figures 1A and 1B, respectively (node locations visualized with the BrainNet Viewer, Xia et al., 2013, http://www.nitrc.org/projects/bnv/). The MNI coordinates are listed in table 2.

**Table 2.**
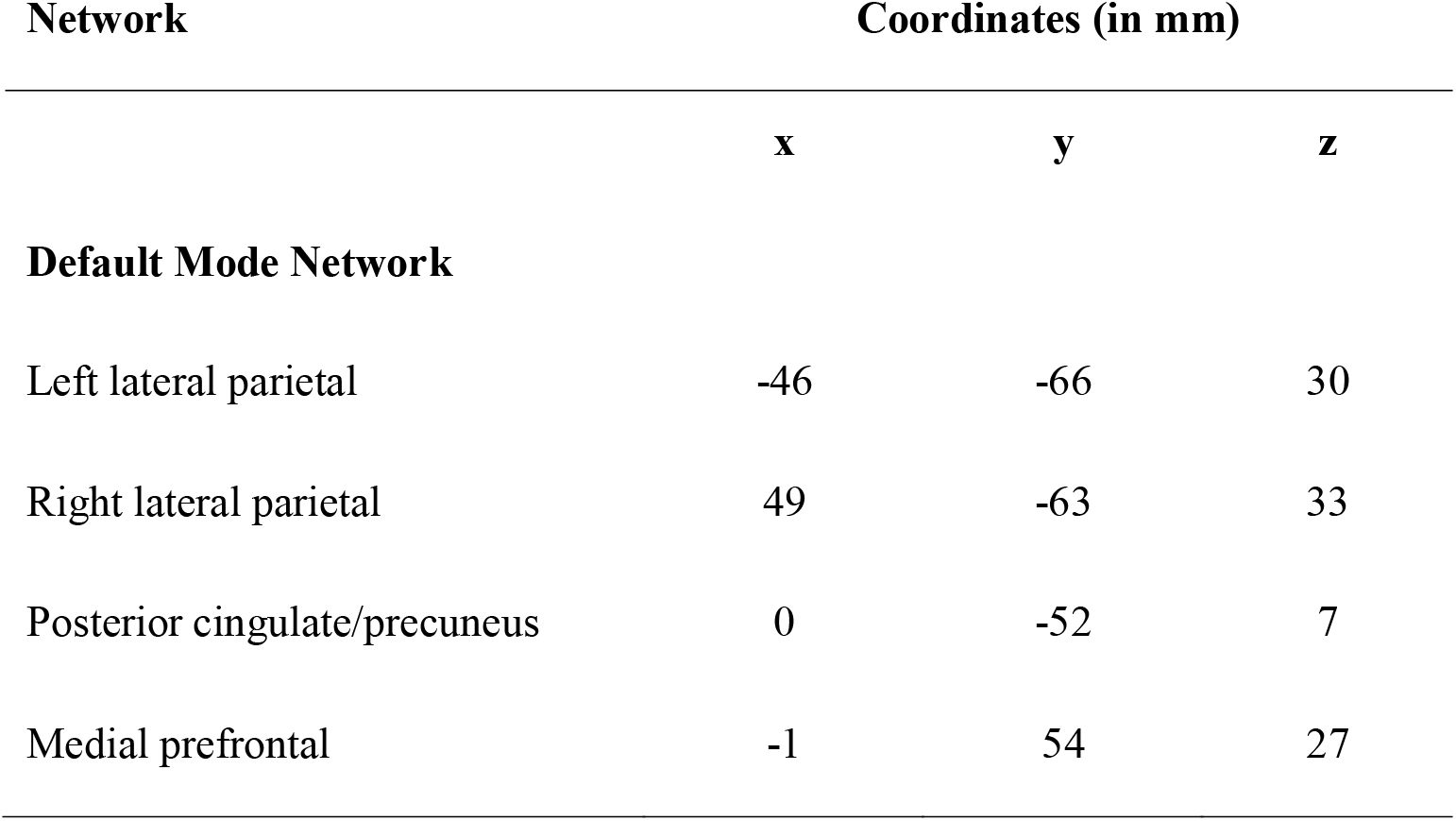
The default mode network nodes and their corresponding MNI coordinates (adapted from Razi et al., 2017).

| Network | Coordinates (in mm) |  |  |
| --- | --- | --- | --- |
|  | x | y | z |
| **Default Mode Network** |  |  |  |
| Left lateral parietal | -46 | -66 | 30 |
| Right lateral parietal | 49 | -63 | 33 |
| Posterior cingulate/precuneus | 0 | -52 | 7 |
| Medial prefrontal | -1 | 54 | 27 |

As shown in figure 1A, the nodes in the DMN were connected via forward, backward, and lateral connections as described in David and collaborators (2006; 2005). Thus, each node was modelled as a point source with the neuronal activity being controlled by operations following the Jansen-Rit model (Jansen & Rit, 1995). These three different types of connections in each model were specified in what is referred in the DCM literature as the ‘A-matrix’. This fully connected model was then estimated for each subject using the DCM for CSD (Friston et al., 2012; Moran et al., 2009; see Supplementary materials for details).

### 4.4 Parametric empirical Bayes

In DCM, the posterior density over the parameters given by the model inversion process is approximated via a variational Bayesian scheme by maximizing a lower bound (the negative free energy) on the log-evidence (Variational Laplace; Friston et al., 2007). A more recent addition, the PEB framework, can be utilized to infer, for example, the group-level commonalities and differences (Friston et al., 2016).

In PEB, the subject-specific parameters – here, the effective connectivity modulations between nodes in DMN – are taken to the group-level and modelled using a General Linear Model (GLM). In doing so, PEB partitions the between-subject variability into designed effects and unexplained random effects (captured by the covariance component). As a special case of Bayesian model selection (BMS), Bayesian model reduction (BMR) enables the inversion of multiple nested models from a single, fully connected (‘full’) model in a hierarchical manner. In doing so it enables a greedy search to compare the negative free energies for the nested models (reduced models), iteratively discarding the parameters that do not contribute to the free energy (originally ‘post-hoc DCM analysis’; Friston & Penny, 2011; Rosa et al., 2012). Consequently, PEB conveys both the estimated group-level connection strengths and their respective uncertainty (posterior covariance component). As such, it is argued that hypotheses about commonalities and differences across subjects can be tested with more precise parameter estimates than with traditional frequentist comparisons (Friston et al., 2016).

A Bayesian Model Average (BMA) is calculated over the best 256 models weighted by their model evidence; for every connection, a posterior probability for the connection being present vs. absent is calculated by comparing evidence from all the models in which the parameter is switched on vs. all the models in which it is switched off. Here, we applied a threshold of >.99 posterior probability, in other words, connections with over .99 posterior probability were retained. The overall process is shown in Figure 2.

### 4.5 Leave-one-out cross-validation

To validate our modelling framework, we investigated which DMN connections are predictive of the subject group by adapting a standard approach in computational statistics, leave-one-subject-out cross-validation (LOSOCV; spm_dcm_loo.m). Here, we iteratively fitted a multivariate linear model (as described in detail in Friston et al., 2016) to provide the posterior predictive density over connectivity changes, which was then used to evaluate the posterior belief of the explanatory variable for the left-out participant: in the present case, the probability of the subject group membership.

To cross-validate a fitted DCM model, one participant was left out each time *before* conducting PEB analysis on the training dataset, and the optimized empirical priors were then used to predict the subject-group to which the dataset from the left-out participant belonged (see Friston et al., 2016 for details). We repeated this procedure for each participant, and in doing so generated probabilities of state affiliation (here, posterior probabilities for subject group-membership).

It is worthwhile to note, that we have estimated the DCM models using the default parameter settings recommended in the literature (Ashburner et al., 2017; Friston et al., 2003; Friston et al., 2012; Kiebel et al., 2009). This is also true for the LOSOCV procedure: no hyper parameter optimization was done. Here, we trained the model with the data from all but the left-out participant (training set), and predicted the state based on the data from the left-out participant (test set) and repeated this procedure leaving out a different participant each time.

### 4.6 Leave-one-subject-out cross-validation

We first estimated predictive performance in a leave-one-subject-out cross-validation paradigm in which LOSOCV metrics for all connections in the DMN and for a hypothesis-driven subsets were estimated (frontoparietal and parietal subsets; figure 3). Next, a data-driven approach was used in which we started the estimation from the connection associated with the largest connectivity reduction between the subject-groups and repeated the procedure for all connections. Here, we utilized a forward stepwise regression in which we started the estimation from the connections with the largest changes and continued through the parameters based on their respective modulation effect sizes. Lastly, we combined connections into data-driven subsets, starting from the connections with the best classification performance, until the classification accuracy stopped improving. The rationale was to investigate the location and direction of the most consistent inter-subject-level effects, in addition to the largest effect sizes identified by the PEB analysis.

### 4.7 Leave-one-state-out cross-validation

Finally, the validation process was generalized by introducing two more difficult classification problems: first, we trained the model on the DCM parameters from the control and the UWS PET-groups, and then tested it on unseen data collected from the MCS* patient-group. Second, we trained the model on the data from the MCS+ and the UWS PET-groups, and again tested on the MCS* datasets. Here, the model was trained on all training datasets. As above, the model used the optimized empirical priors to predict the more likely patient-group the test dataset (MCS*) belonged. We hypothesized that if our modelled effects are valid, and if the sustained PET-metabolism reflects higher level of consciousness present in the MCS* patients in comparison to UWS PET-patients, in the former case the model should classify the test datasets as controls rather than UWS PET-. Similarly, in the latter case, given that the MCS+ patients are conscious, the test data should be classified as MCS+ rather than UWS PET-. Here, we used posterior probability for subject group-membership to quantify classification performance.

## Acknowledgements and financial disclosure

We gratefully acknowledge support from the University of Kent’s School of Computing and GIGA Doctoral School for Health Sciences. The study was further supported by the University and University Hospital of Liège, the Belgian National Funds for Scientific Research (FRS-FNRS), the European Union’s Horizon 2020 Framework Programme for Research and Innovation under the Specific Grant Agreement No. 945539 (Human Brain Project SGA3), the FNRS MIS project (F.4521.23), the FNRS PDR project (T.0134.21), the ERA-Net FLAG-ERA JTC2021 project ModelDXConsciousness (Human Brain Project Partnering Project), the fund Generet, the King Baudouin Foundation, the Télévie Foundation, the European Space Agency (ESA) and the Belgian Federal Science Policy Office (BELSPO) in the framework of the PRODEX Programme, the BIAL Foundation, the Mind Science Foundation, the European Commission, and the Fondation Leon Fredericq.

## 7 Supporting Materials

**Table S1.**
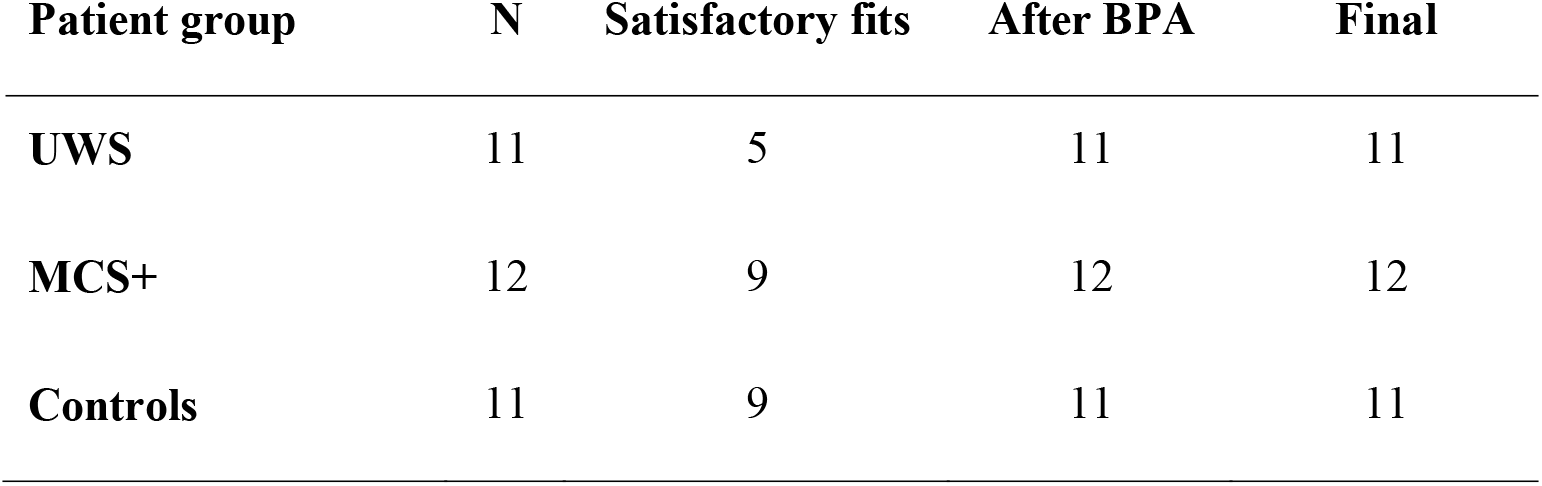
The number of satisfactory fits with the default hyperparameters and after adjusting the neural innovations and the noise precision for the different subject groups.

**Figure S1.**
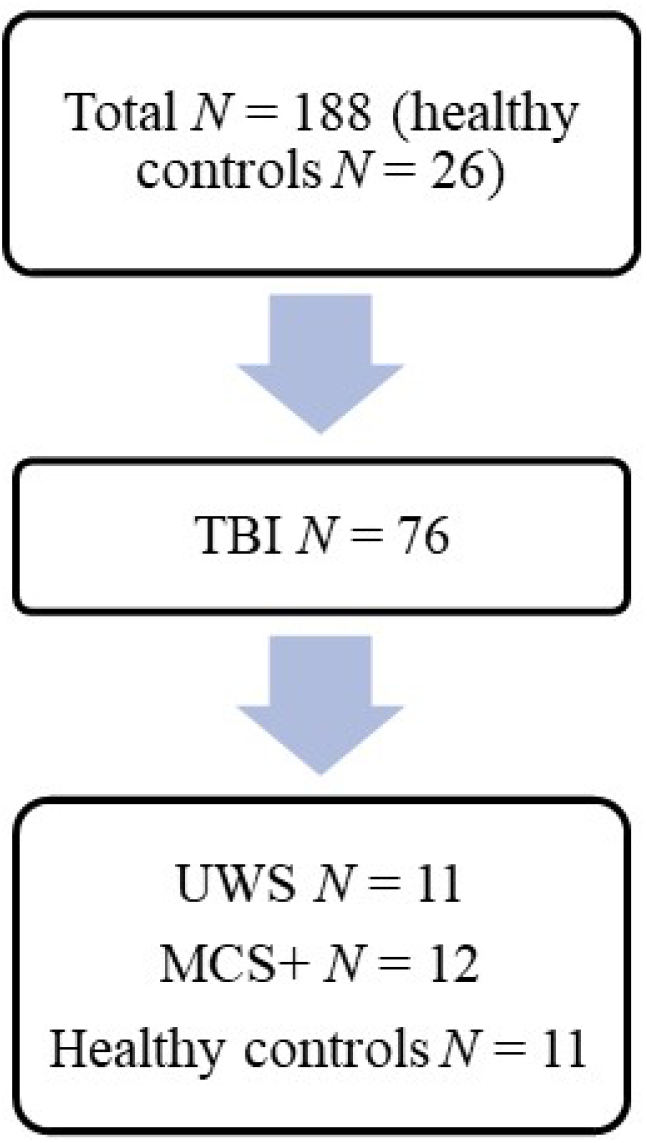
A flowchart showing the dataset pruning process, and the corresponding *N* for the experimental groups. From the full dataset, patients with TBI (*N* = 76) were identified. Next, the main group of interest – patients diagnosed as UWS (*N* = 11) – were distinguished. We then pseudo-randomly drew a cohort of 11 healthy controls to adjust for the group-size discrepancies. A cohort of 12 MCS+ patients were identified to act as a second control group.

**Figure S2.**
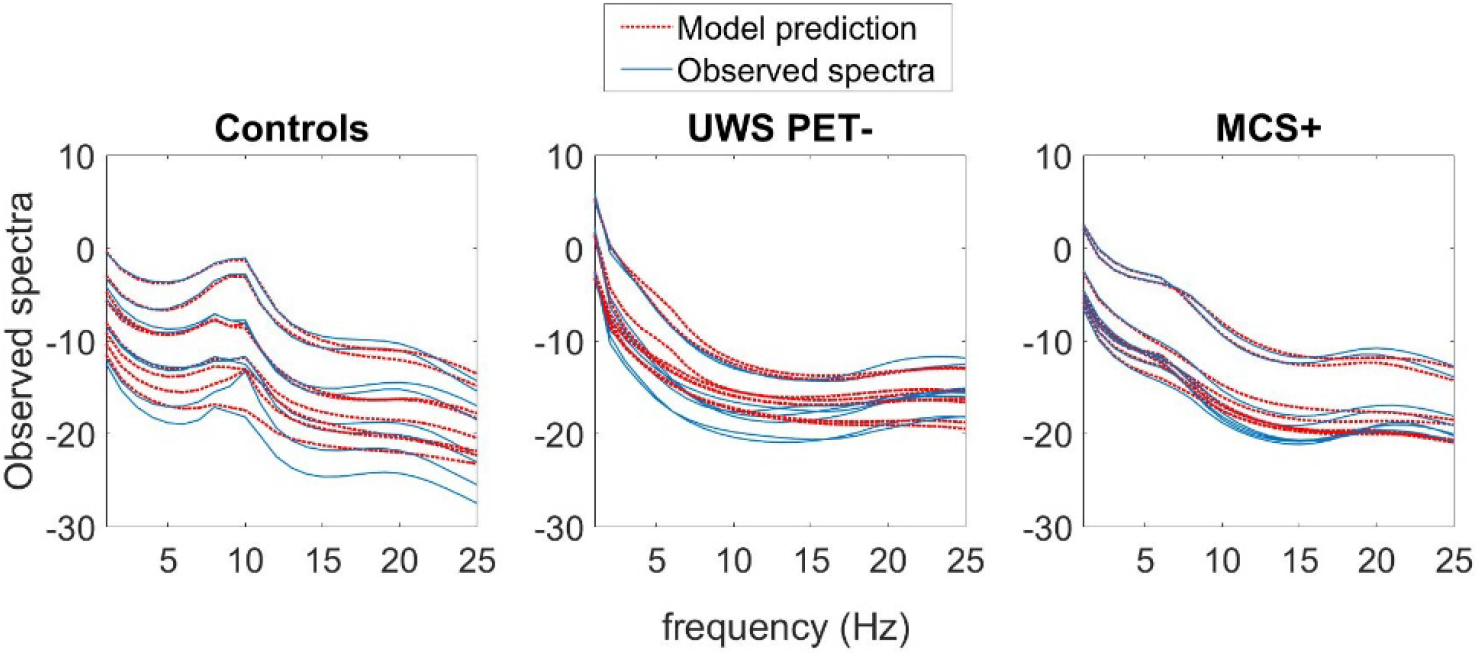
The average model fits across the participants in all subject-groups. A-C. Subject-averaged power spectra of the observed EEG channel-space data, juxtaposed with that predicted by the fitted DCM models of each subject group. Individual lines reflect spatial modes.

**Figure S3.**
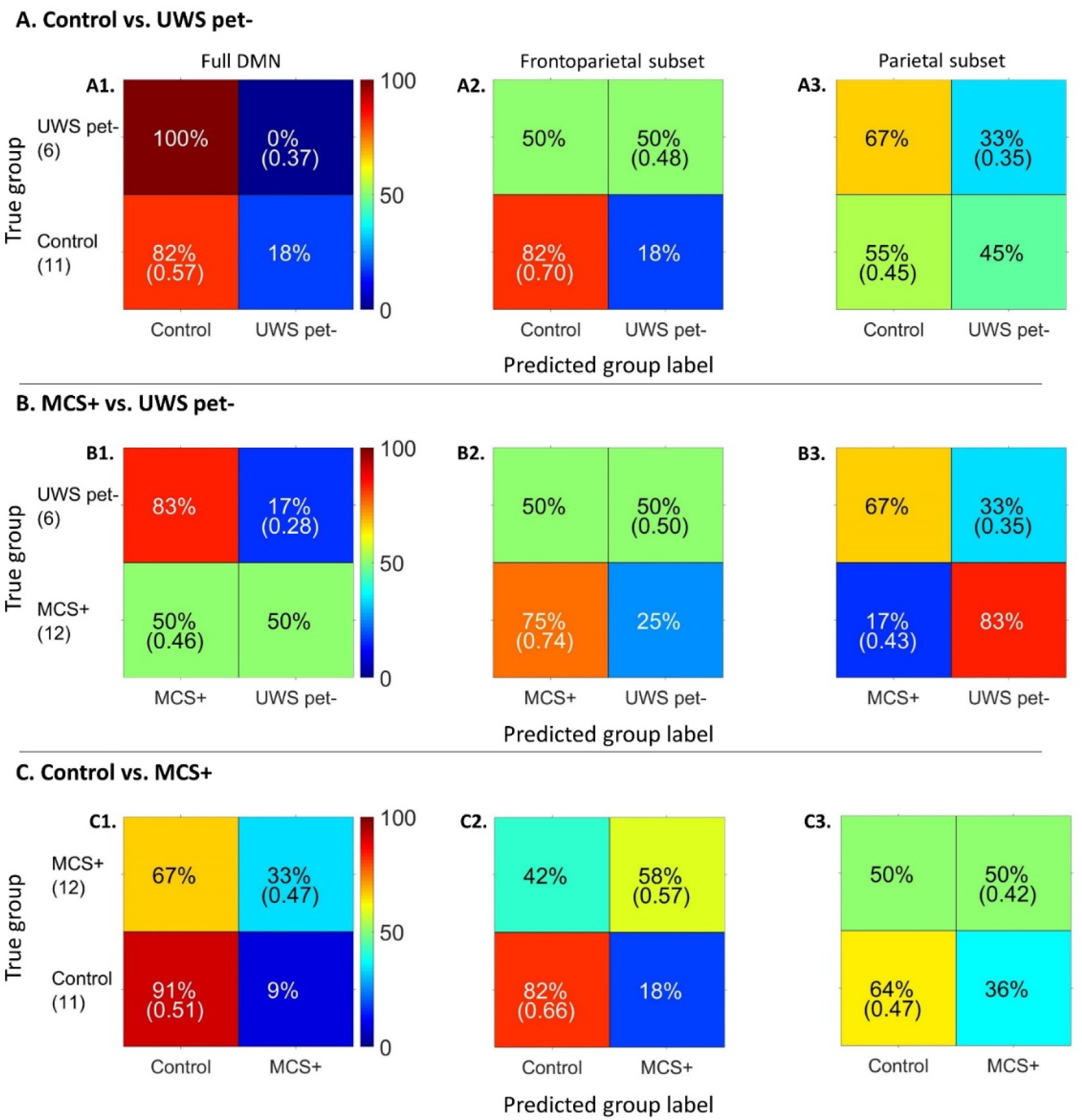
Classification accuracy percentage (mean posterior probability for correct classification) in the leave-one-subject-out cross-validation paradigm for the hypothesis-driven subsets. The number of subjects in each group is shown in parenthesis under the true group labels. The frontoparietal subset performed the best in terms of both classification accuracy and mean posterior probability, especially with healthy controls for healthy control vs. UWS PET– and MCS+ vs. UWS PET-contrasts (panels A2 and B2, respectively). Classification based on full DMN had high accuracy for healthy controls; however, the mean posterior probabilities bordered the chance level.

**Figure S4.**
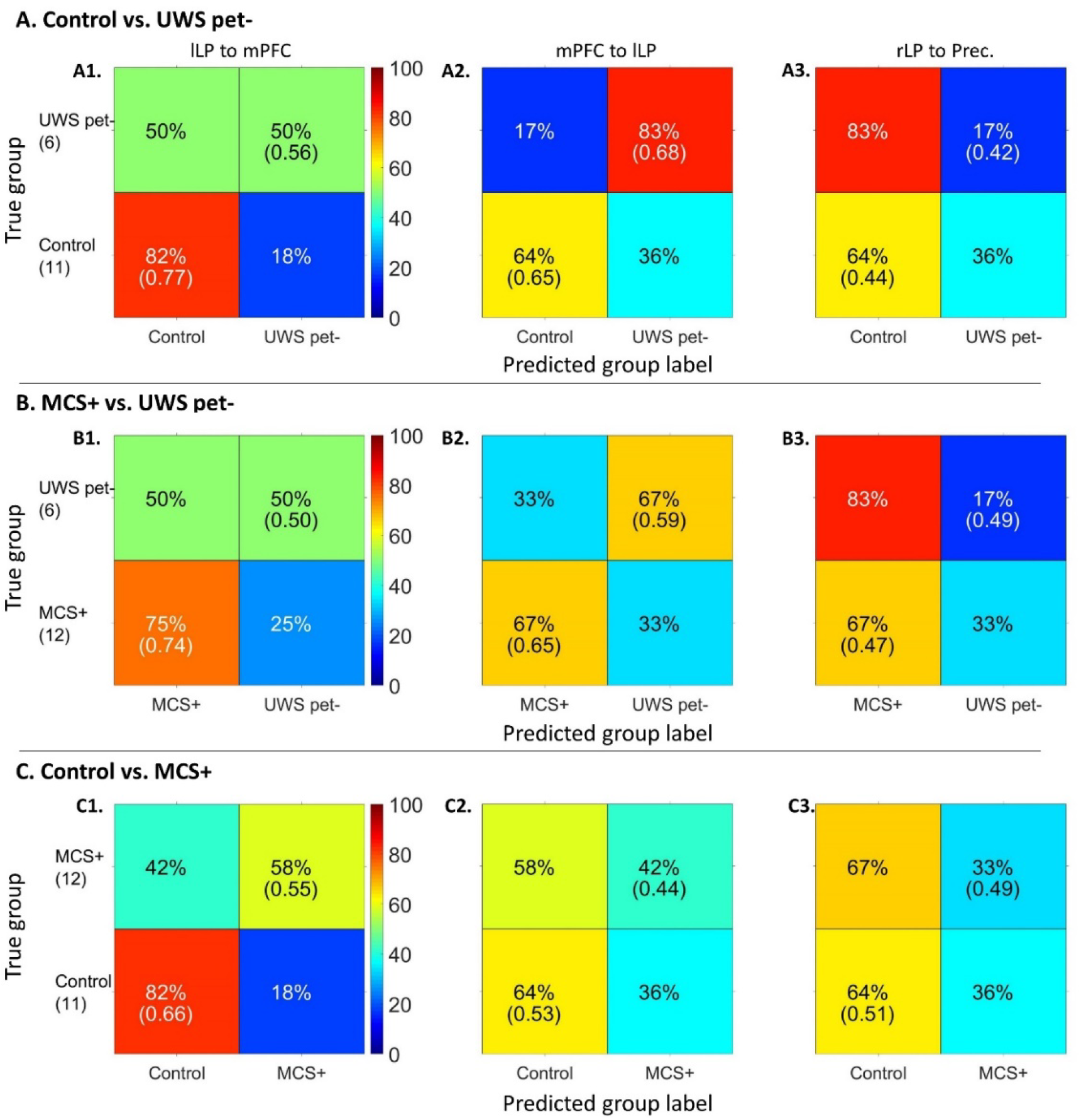
Classification accuracy percentage (mean posterior probability for correct classification) in the leave-one-subject-out cross-validation paradigm for the data-driven approach. The number of subjects in each group is shown in parenthesis under the true group labels. For the healthy controls vs. UWS PET– and MCS+ vs. UWS PET-contrasts, the frontoparietal backward connection from mPFC to lLP performed best in terms of both classification accuracy and mean posterior probability. Forward frontoparietal connectivity from lLP to mPFC classified healthy controls and MCS+ patients from UWS PET-with high accuracy but bordered the chance level with UWS PET-. Similarly, lLP to mPFC connectivity performed the best with the healthy controls vs. MCS+ contrast.

