## Supplementary materials for "Lateral frontoparietal effective connectivity differentiates and predicts state of consciousness in traumatic disorders of consciousness"

### 1 Supplementary Materials

#### 1.1 Data acquisition

The dataset consisted of the patient data with 26 healthy controls (total *N* = 188). TBI, as compared to non-traumatic aetiologies has been associated with more focal injury centred often on areas susceptible to rotational forces, such as the brainstem, midbrain, thalamus, hypothalamus, cerebellum, and posterior corpus callosum (Guldenmund et al., 2016; Newcombe et al., 2010). Hence, to lower the chance of modelling heavily damaged areas, we first identified patients with TBI (*N* = 76). Patients admitted due to any other aetiology, e.g. anoxia or haemorrhage, were thus excluded from the further analyses. Next, among the TBI patients, we identified the main group of interest, i.e. patients diagnosed with UWS (Laureys et al., 2010, *N* = 11). The UWS patients were further divided into two groups based on their PET metabolism (see next section). Using the random number generator in MATLAB, we then pseudo-randomly drew a cohort of 11 healthy control subjects to adjust for the group-size discrepancies. From the TBI dataset, we furthermore identified a cohort of 12 MCS+ patients, to act as a second control group. MCS+ patients are characterised by fluctuating but reproducible signs of consciousness with partially preserved language related behaviours (Aubinet et al., 2020; Bruno et al., 2012; Thibaut et al., 2020). We rationalised that by including a group of conscious patients to contrast with UWS patients, who nonetheless suffer from traumatic brain injury, the probability for the results reflecting only brain damage – and not consciousness itself – decreases. Figure s1 shows a flowchart for the dataset pruning process.

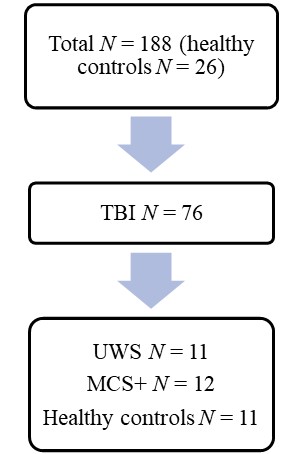

**Figure s1.** A flowchart showing the dataset pruning process, and the corresponding *N* for the experimental groups. From the full dataset, patients with TBI (*N* = 76) were identified. Next, the main group of interest – patients diagnosed as UWS (*N* = 11) – were distinguished. We then pseudo-randomly drew a cohort of 11 healthy controls to adjust for the group-size discrepancies. A cohort of 12 MCS+ patients were identified to act as a second control group.

#### 1.2 Behavioural and Positron Emission Tomography assessments

Patients were behaviourally assessed on the day of the PET and EEG imaging using the Coma Recovery Scale – Revised (CRS-R; Kalmar & Giacino, 2005b) five to seven times a day, and the diagnosis was based on the highest score obtained.

Positron Emission Tomography Data Collection and Analysis Flurodesoxyglucose-PET (FDG-PET) scans were acquired from all the included patients using the methodology as described in (Stender et al., 2014a). The scans were acquired on a Philips Gemini TF PET-CT scanner (Philips Medical Systems) approximately 30 minutes after an intravenous injection of 150 or 300 MBq of the radioactive tracer, fluor-18 fluordeoxyglucose (FDG). The brain imaging was obtained during an awake-period and while eyes open in a silent and dark room (ensured by an examiner present by administering tactile or auditory stimuli when the patients were closing their eyes). The data analysis identified relatively preserved and decreased metabolism in patients in comparison to controls and was conducted using Statistical Parametric Mapping (SPM8).

The analysis results were visually inspected by a trained neurologist to reach a PET+- or PET- -diagnosis following previous findings. A PET--diagnosis was produced by a complete bilateral hypometabolism of the associative frontoparietal cortex with no voxels with preserved metabolism, whereas PET+-diagnosis was produced by an incomplete hypometabolism and partial preservation of activity within these areas (Laureys et al., 2004; Nakayama et al., 2006; Thibaut et al., 2012).

#### 1.3 Dynamic Causal Modeling: Model inversion

In DCM, model inversion refers to the process of fitting a model to explain the empirical data of each participant’s dataset, and thereby inferring a full probability density over the possible values of model parameters (with the expected values and covariance). The default parameter settings in DCM for CSD led to inaccurate fits of the model when inspected visually (see van Wijk et al., 2018, *p.* 824). To address this, we doubled the number of maximum iterations to 256 and estimated the models with two adjustments to the hyperparameters: first, the shape of the neural innovations (i.e. baseline neural activity) were set to flat (-32) instead of the default white and pink (1/f) noise-component mixture (Moran et al., 2009). Second, we increased the noise precision value from 8 to 12 to bias the inversion process towards accuracy over complexity (see Friston et al., 2012 and Moran et al., 2009). With these adjustments, we estimated the full DMNs again, and applied the Bayesian Parameter Averaging (BPA) for each of the subject-groups separately, averaging over the posterior fits from the subjects for whom the model did converge satisfactory and setting these averaged posteriors as new priors for the respective non-converged subjects. Finally, we estimated all the full models again for all the subjects with setting the posteriors from the earlier subject model estimations as updated priors, but this time with the neural innovations and noise precision set back to default settings. To validate that the priors we used in the final inversion were suitable, we compared the group-level model evidence obtained from the BMA with and without the adjusted noise levels. With all comparisons, the default hyperparameter settings with the updated priors generated better model evidence (difference in free energies for control vs. UWS PET-, control vs. MCS+, and MCS+ vs. UWS PET- were +29364, +3096, and +39726, respectively. To qualitatively assess the model fits, the observed and model-predicted cross-spectra were visually compared in each participant and judged sufficiently similar. This process led to satisfactory fits for the subjects (table s1). The average fits over subjects for each group are shown in Figure s2.

**Table s1.** The number of satisfactory fits with the default hyperparameters and after adjusting the neural innovations and the noise precision for the different subject groups.

| **Patient group** | **N** | **Satisfactory fits** | **After BPA** | **Final** |
| --- | --- | --- | --- | --- |
| **UWS** | 11 | 5 | 11 | 11 |
| **MCS+** | 12 | 9 | 12 | 12 |
| **Controls** | 11 | 9 | 11 | 11 |

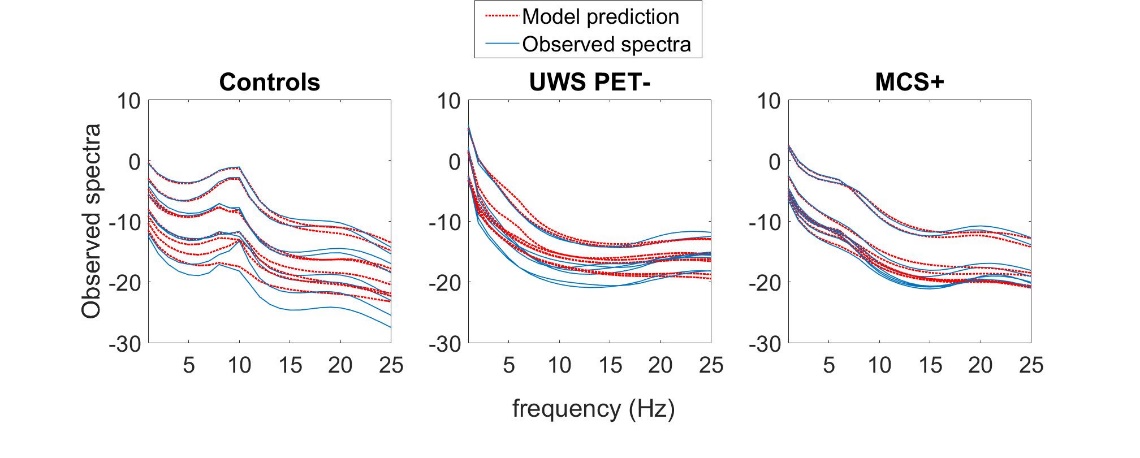

**Figure s2.** The average model fits across the participants in all subject-groups. A-C. Subject-averaged power spectra of the observed EEG channel-space data, juxtaposed with that predicted by the fitted DCM models of each subject group. Individual lines reflect spatial modes.

#### 1.4 Results

Figures s2 and s3 show confusion matrices of prediction accuracy calculated by labelling posterior probabilities greater than 0.5 as a positive classification, for the hypothesis-driven subsets and the data-driven approach, respectively. In figure s3, the frontoparietal subset performed most consistently in terms of both, classification accuracy and mean posterior probability, especially with healthy controls.

In figure s4 with the data-driven approach, the frontoparietal backward connection from mPFC to lLP performed best in terms of both, classification accuracy and mean posterior probability for healthy control vs. UWS PET- and MCS+ vs. UWS PET- contrasts. Forward frontoparietal connectivity from lLP to mPFC classified healthy controls and MCS+ patients from UWS PET- with high accuracy but bordered the chance level with UWS PET-. Overall, the bi-directional left frontoparietal connections provided the best classification performances amongst all connections and combinations tested.

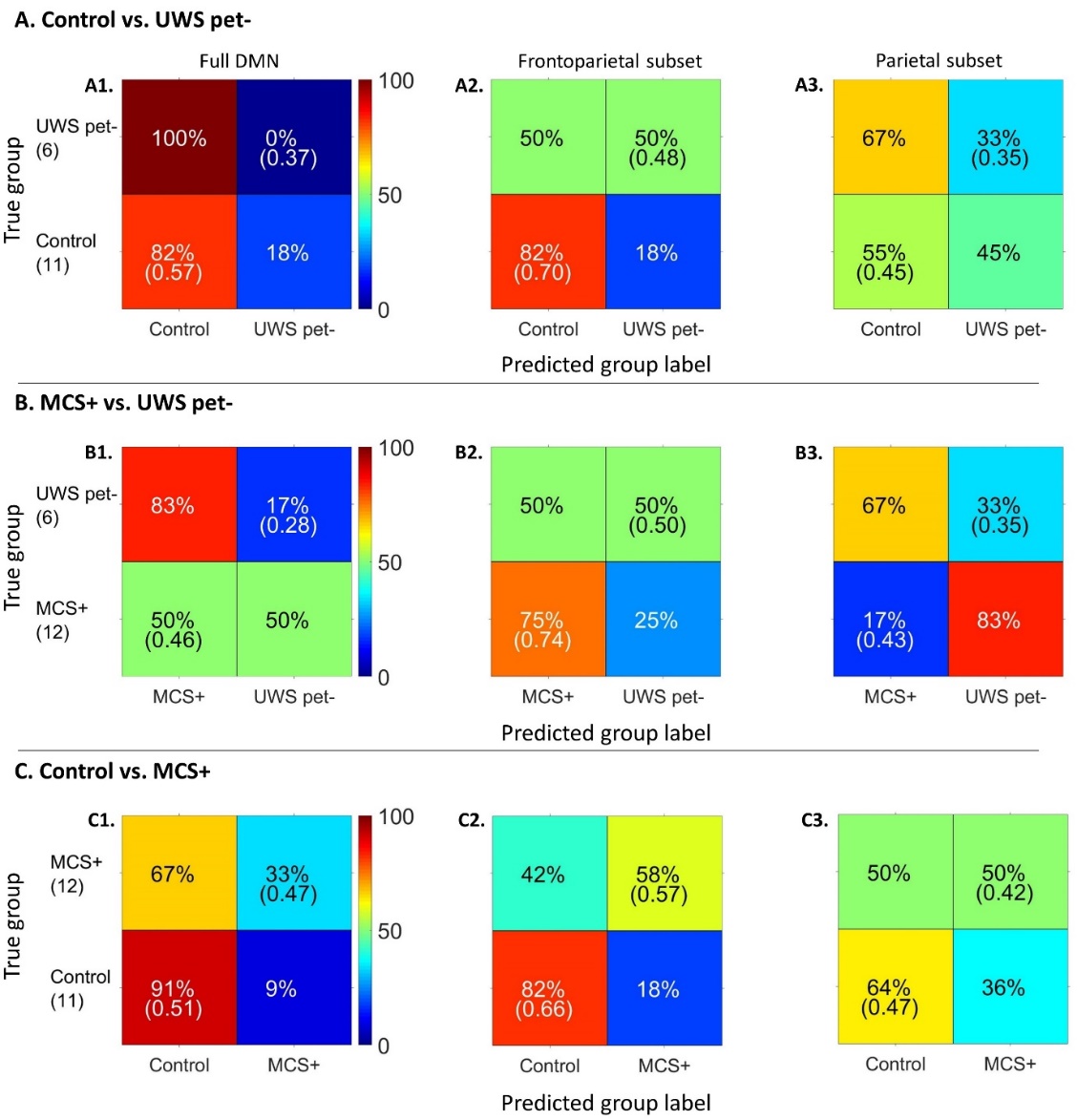

**Figure s3.** Classification accuracy percentage (mean posterior probability for correct classification) in the leave-one-subject-out cross-validation paradigm for the hypothesis-driven subsets. The number of subjects in each group is shown in parenthesis under the true group labels. The frontoparietal subset performed the best in terms of both classification accuracy and mean posterior probability, especially with healthy controls for healthy control vs. UWS PET- and MCS+ vs. UWS PET- contrasts (panels A2 and B2, respectively). Classification based on full DMN had high accuracy for healthy controls; however, the mean posterior probabilities bordered the chance level.

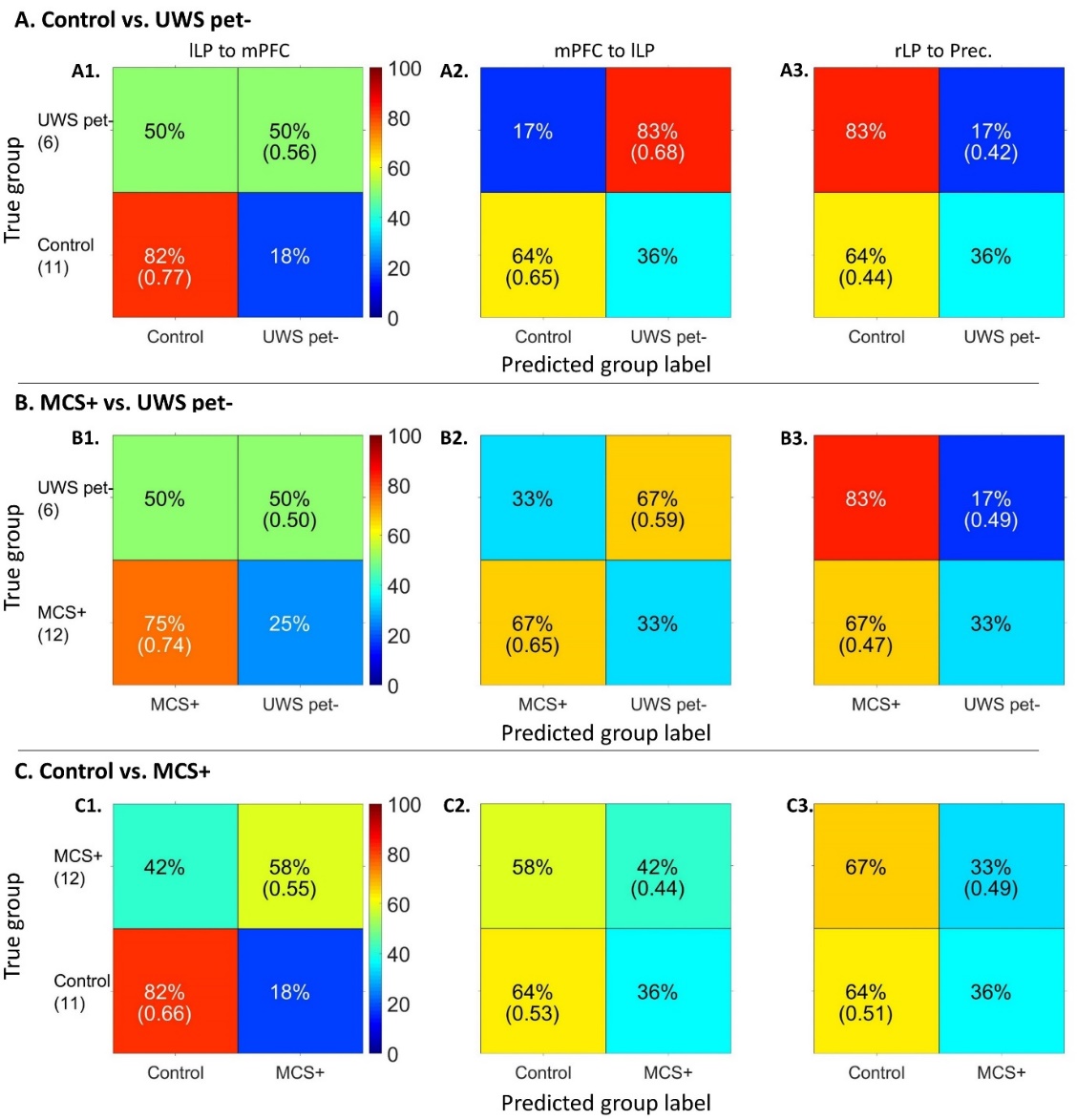

**Figure s4.** Classification accuracy percentage (mean posterior probability for correct classification) in the leave-one-subject-out cross-validation paradigm for the data-driven approach. The number of subjects in each group is shown in parenthesis under the true group labels. For the healthy controls vs. UWS PET- and MCS+ vs. UWS PET- contrasts, the frontoparietal backward connection from mPFC to lLP performed best in terms of both, classification accuracy and mean posterior probability. Forward frontoparietal connectivity from lLP to mPFC classified healthy controls and MCS+ patients from UWS PET- with high accuracy but bordered the chance level with UWS PET-. Similarly, lLP to mPFC connectivity performed the best with the healthy controls vs. MCS+ contrast.
